# Chemical inhibition of ENL/AF9 YEATS domains in acute leukemia

**DOI:** 10.1101/2020.12.01.406694

**Authors:** Leopold Garnar-Wortzel, Timothy R. Bishop, Seiya Kitamura, Natalia Milosevich, Joshua N. Asiaban, Xiaoyu Zhang, Qinheng Zheng, Emily Chen, Anissa R. Ramos, Christopher J. Ackerman, Eric N. Hampton, Arnab K. Chatterjee, Travis S. Young, Mitchell V. Hull, K. Barry Sharpless, Benjamin F. Cravatt, Dennis W. Wolan, Michael A. Erb

**Affiliations:** Department of Chemistry, Scripps Research, La Jolla, CA, USA; Department of Molecular Medicine, Scripps Research, La Jolla, CA, USA; Department of Integrative Structural and Computational Biology, The Scripps Research Institute, La Jolla, California, USA; California Institute for Biomedical Research, Scripps Research, La Jolla, CA, USA

## Abstract

Transcriptional co-regulators, which mediate chromatin-dependent transcriptional signaling, represent tractable targets to modulate tumorigenic gene expression programs with small molecules. Genetic loss-of-function studies have recently implicated the transcriptional co-activator, ENL, as a selective requirement for the survival of acute leukemia and highlighted an essential role for its chromatin reader YEATS domain. Motivated by these discoveries, we executed a screen of nearly 300,000 small molecules and identified an amido-imidazopyridine inhibitor of the ENL YEATS domain (IC_50_ = 7 µM). Leveraging a SuFEx-based high-throughput approach to medicinal chemistry optimization, we discovered SR-0813 (IC_50_ = 25 nM), a potent and selective ENL/AF9 YEATS domain inhibitor that exclusively inhibits the growth of ENL-dependent leukemia cell lines. Armed with this tool and a first-in-class ENL PROTAC, SR-1114, we detailed the response of AML cells to pharmacological ENL disruption for the first time. Most notably, displacement of ENL from chromatin by SR-0813 elicited a strikingly selective suppression of ENL target genes, including *HOXA9/10, MYB, MYC* and a number of other leukemia proto-oncogenes. Our study reproduces a number of key observations previously made by CRISPR/Cas9 loss of function and dTAG-mediated degradation, and therefore, both reinforces ENL as an emerging leukemia target and validates SR-0813 as a high-quality chemical probe.

## Introduction

Transcriptional dysregulation in cancer pathogenesis represents a potentially rich source of targets for therapeutic intervention. In AML, genetic alterations disproportionately affect transcriptional machinery and frequently converge on similar nodes of pathogenic gene control, like the activation of *HOXA* cluster genes.^1–9^ Tragically, despite being the second-most-common leukemia, the 5-year survival rate of patients with AML remains around 25%. Therefore, there is significant interest in drugs and drug targets that would enable the core transcriptional circuitries driving AML pathogenesis to be suppressed. Motivated to illuminate such targets, we previously applied genome-scale CRISPR/Cas9 loss-of-function screening to identify genes that are required for AML survival.^10^ Through this work, and contemporaneously with others, we identified the transcriptional co-activator, ENL (encoded by *MLLT1*), as a requirement for the proliferation of AML and B-ALL (B-cell acute lymphoblastic leukemia) that harbor oncogenic *multiple lineage leukemia* (*MLL*) rearrangements.^10,11^ Since it is not required for the survival of normal hematopoietic stem and progenitor cells (HSPCs) *ex vivo*,^10,11^ we proposed ENL as an attractive target for the development of anti-leukemia therapeutics.

ENL is a chromatin reader protein possessing an amino-terminal YEATS domain (named for the first-discovered members of the family: Yaf9, ENL, AF9, Taf14, Sas5) and a disordered carboxy-terminal protein-protein interaction (PPI) interface.^12^ Adapted to bind acylated lysine side chains, the YEATS domain of ENL mediates its localization to acetylated regions of chromatin, including enhancers and promoters.^10,11^ The localization of ENL to these *cis*-regulatory elements then allows for the assembly of additional transcriptional effectors – such as the super elongation complex (SEC) – that regulate transcription by RNA polymerase (Pol) II.^10,11,13^ Presently, the ENL YEATS domain is known to bind with low micromolar affinity to acetyl (ac) and crotonyl (cr) modifications of H3K9 (histone H3 Lys9), H3K18, and H3K27.^10,11,14^ While these post-translational modifications are dispersed throughout *cis*-regulatory elements of the genome, ENL is asymmetrically distributed to its binding sites in AML cells.^10^ That is, a small number of genes, including a disproportionate number of leukemia proto-oncogenes and dependencies, are occupied with a disproportionate amount of ENL at their promoters. To understand the impact of this asymmetry on ENL-dependent transcriptional control, we previously used degradation tag (dTAG) technology to engineer a tagged ENL allele that is sensitive to pharmacologically induced degradation.^10,15^ The rapid kinetics of this system allowed us to probe the direct effects of ENL loss at early time points and revealed that asymmetrically loaded ENL target genes are preferentially sensitive to loss of ENL. As a result, a number of disease-relevant leukemia drivers, such as *HOXA9*/*10, MEIS1, MYB*, and *MYC*, are selectively suppressed by loss of ENL.^10^

The YEATS domain represents an attractive target to disrupt ENL function with small molecules as it is: (i) required to mediate stable association of ENL with chromatin and (ii) essential for the proliferation of ENL-dependent leukemia.^10,11^ Moreover, recent disclosures of small-molecules and synthetic peptide ligands have convincingly established the druggability of ENL and AF9 (encoded by *MLLT3*) YEATS domains.^16–21^ However, the phenotypes observed in AML cells following genetic suppression of ENL have yet to be validated with direct-acting chemical tools. The most potent small-molecule reported to date, SGC-iMLLT (*K*_d_ = 129 nM), has not been studied in AML cells beyond proximal measures of target engagement,^17^ and a previously reported synthetic peptide inhibitor, XL-13m, does not feature sufficient intracellular activity to efficiently displace ENL from chromatin.^19^ In each case, only a handful of ENL target genes have been studied to confirm on-target effects of the ligands and no impact on ENL-dependent leukemia proliferation has been reported. Therefore, pharmacological validation of ENL as a therapeutic target remains unexplored, underscoring the need to discover and fully characterize potent and selective ENL chemical probes.

Here, we report on the results of our ongoing efforts to develop advanced chemical tools able to probe the cellular function of ENL in physiology and disease. Through a coordinated effort in discovery chemistry that began with high-throughput chemical screening, we developed an ENL degrader, SR-1114, and ENL YEATS inhibitor, SR-0813 (*K*_d_ = 30 nM). Our discovery of SR-0813 relied on a high-throughput medicinal chemistry approach that leverages biocompatible SuFEx transformations to synthesize hit analogs in highly parallel, miniaturized formats. As previously demonstrated, the crude products of these reactions can then be tested directly in biochemical assays without further purification, expediting the process of medicinal chemistry.^22^ Here, we extend this approach by demonstrating that SuFEx-based high-throughput medicinal chemistry is compatible with cell-based measures of activity, ultimately enabling the identification of SR-0813. Mechanistically studied in AML model systems with integrative transcriptional genomics, we define the impact of SR-0813 on pathogenic gene control and highlight the remarkably selective suppression of leukemic drivers. Together, these studies provide the first pharmacological validation of ENL as an anti-leukemia target and advance a high-quality chemical probe to study ENL/AF9 YEATS domains.

## Results

In prior research efforts, we and others used mutational approaches to identify a critical role for the YEATS domain in ENL-dependent leukemia proliferation.^10,11^ Interested to validate these results with direct-acting pharmacological tools, we undertook efforts to discover chromatin-competitive inhibitors of the ENL YEATS domain. Using a homogenous time-resolved FRET (HTRF)-based measure of binding between the ENL YEATS domain and H3K27cr, we screened for putative ENL YEATS inhibitors from a collection of approximately 275,000 diverse small molecules (Figure 1a). Confirmed primary hits were tested in dose-response format, leading to the selection of 100 compounds for hit-expansion studies and further evaluation. In total, these 100 hits and an additional 275 structurally similar compounds were tested in dose-response assays against ENL YEATS and then counter-screened against the first bromodomain (BD1) of BRD4 (a structurally dissimilar protein fold that also recognizes acetyl-lysine residues).^23^ To validate ENL binding by an orthogonal measure and to prioritize cell-permeable compounds, YEATS-selective hits were evaluated for intracellular target engagement using a luminescence-based cellular thermal shift assay (CETSA).^21^ These efforts resulted in the identification of compound **1**, an amido-imidazopyridine inhibitor of the ENL YEATS domain (Figure 1b-d, Figure S1a).

**Figure 1.**
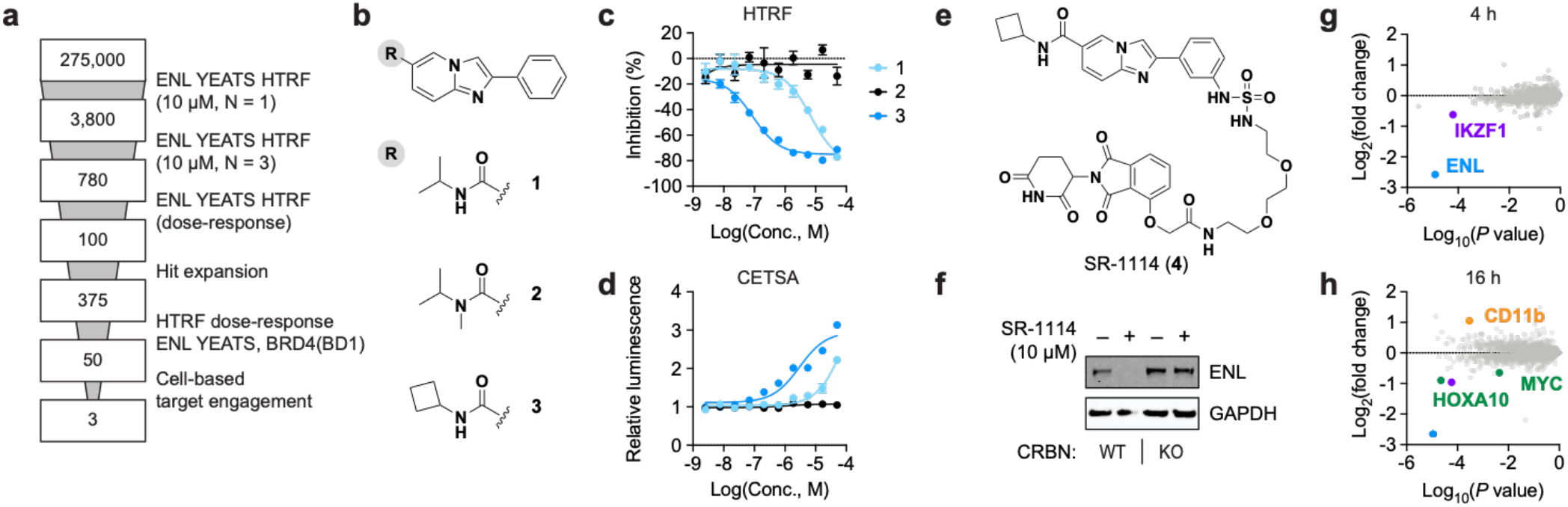
An amido-imidazopyridine scaffold with activity against ENL/AF9 YEATS domains. (a) Summary of screening campaign to identify and validate ligands for the ENL YEATS domain. (b) Structure of screening hit, **1**, and compounds used for preliminary SAR-finding studies. (c) Activity against the ENL YEATS domain measured by HTRF. Mean percent inhibition ± s.e.m., *n* = 4. (d) Engagement of ENL(YEATS)-HiBiT measured by ligand-induced luminescence. Signal normalized to DMSO ± s.e.m., *n* = 4. (e) chemical structure of SR-1114 (**4**). (f) Immunoblot confirmation of CRBN-dependent ENL degradation by SR-1114. (g, h) Changes in protein abundance measured in MV4;11 cells by expression proteomics after 4-h (g) and 16-h (h) treatments of SR-1114 (10 µM). *P* value derived from two-tailed Student’s *t*-test for 6,972 proteins with at least 2 unique spectral counts detected.

To better gauge the suitability of this scaffold for further optimization, we sought to identify preliminary evidence of a structure-activity relationship (SAR). Methylation of the amide (**2**) resulted in complete ablation of activity against ENL YEATS, whereas exchanging the isopropyl for a cyclobutyl group (**3**) greatly improved potency by both HTRF and CETSA (Figure 1b-d, Figure S1a). Limited exploration of the phenyl ring did not yield any notable improvements but did establish the meta and para positions as permissible sites for substitution. We speculated that these sites might tolerate linker attachments for proteolysis targeting chimeras (PROTACs), ultimately considering that the discovery of a selective degrader would substantiate ENL as an authentic target of the imidazopyridine series. Indeed, SR-1114 (**4**), a PROTAC based on attachment of thalidomide to compound **3**, elicits rapid, cereblon (CRBN)-dependent degradation of ENL in AML cells (Figure 1e,f; Figure S1b-d). After a 4-hour exposure of MV4;11 cells to SR-1114, mass spectrometry (MS)-based proteomics revealed a highly selective effect on ENL abundance (Figure 1g). After 16 hours, SR-1114 preferentially suppressed the ENL target genes, HOXA10 and MYC (Figure 1h, Table S1), closely replicating the effects of dTAG-mediated ENL degradation.^10^ Furthermore, SR-1114 treatment increased CD11b abundance, which is in agreement with its proposed role in preventing the terminal differentiation of AML cells.^11^

These data elevated our confidence in ENL as a leukemia target and provided orthogonal validation of ENL as a target of the amido-imidazopyridine series. However, proteomics studies also revealed weak off-target degradation of the IKAROSE family transcription factor, IKZF1 (Figure 1g,h). Degradation of IKZF1 and other CRBN neo-substrates is a recurring liability of CRBN-based PROTACs,^24,25^ which in this case, might complicate the study of ENL-dependent transcriptional control in acute leukemia. Therefore, we pursued enhancements in ligand affinity that would (i) enable study of the YEATS domain in acute leukemia and (ii) provide better starting points for more potent and selective ENL degraders.

Additional improvements to **1** were discovered using a high-throughput hit-to-lead process that leverages sulfur(VI) fluoride exchange (SuFEx) “click” transformations for parallelized chemical diversification (Figure 2a).^22^ Capitalizing on the high-yielding reaction of iminosulfur oxydifluoride SuFEx hubs with primary and secondary amines,^26^ a diverse library of sulfamide- or sulfuramidimidoyl-fluoride-linked analogs can be rapidly assembled in miniaturized formats.^22^ Importantly, the biocompatible nature of these reactions permits testing of crude products directly in biochemical assays without need of further purification.^22^ Without a crystal structure to guide iterative medicinal chemistry efforts, we attempted to expedite empirical discovery of optimized ENL ligands using this approach.

**Figure 2.**
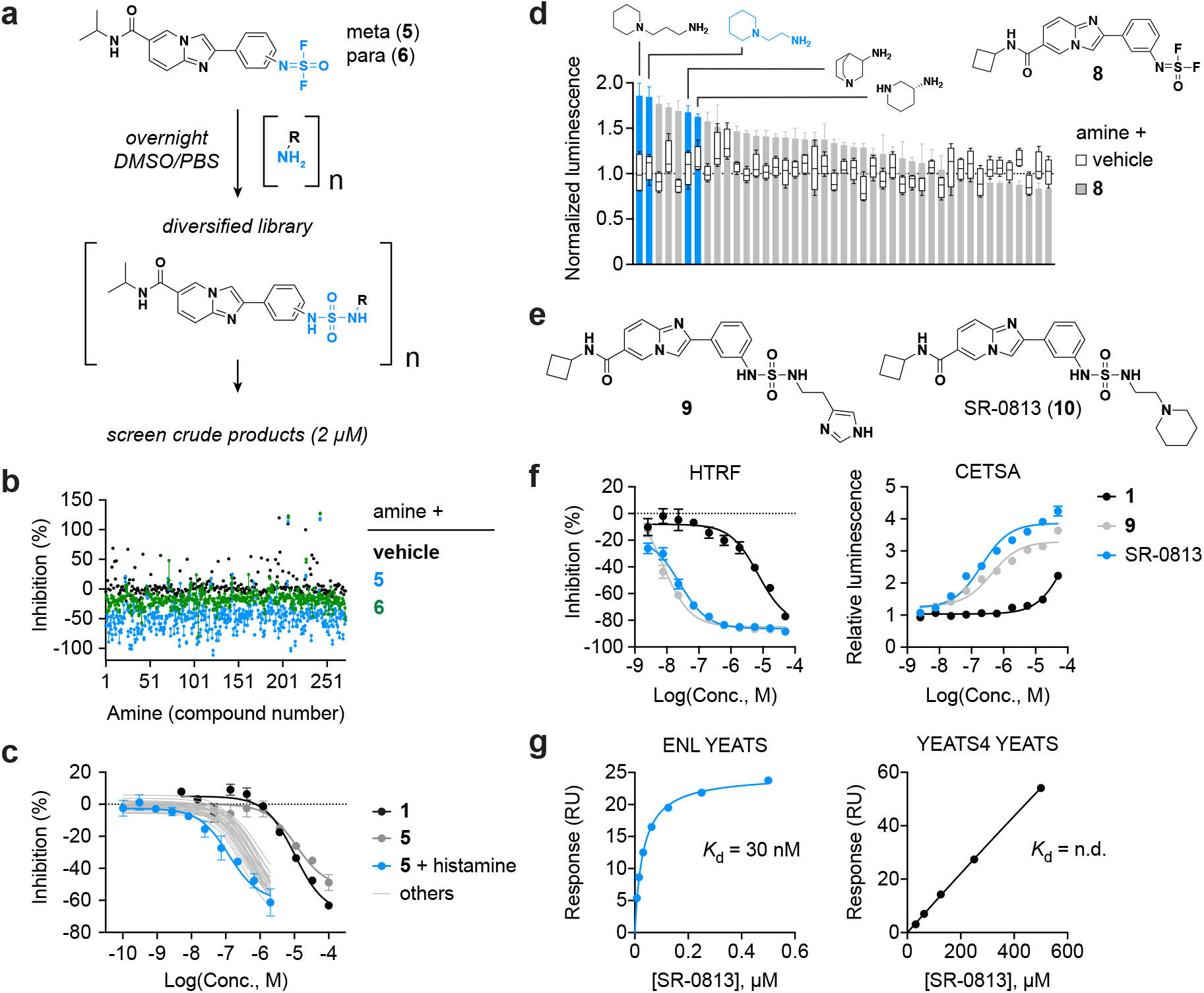
Hit optimization by SuFEx-based high-throughput medicinal chemistry. (a) scheme for highly parallel medicinal chemistry approach. Iminosulfur oxydifluoride SuFEx hubs react with primary (shown) and secondary (not shown) amines to assemble a diverse collection of near derivatives that can be tested as crude reaction products. (b), ENL YEATS HTRF assay to measure activity of 288 amines reacted with vehicle (*n* = 1), **5** (*n* = 2), or **6** (*n* = 2). Individual replicates are connected by a vertical line. (c) Measurement of dose-responsive activity for 28 hits in the screen, testing crude reaction products by ENL YEATS HTRF. Mean ± s.e.m., *n* = 2. (d) Activity of crude SuFEx reaction products measured in living cells by ENL(YEATS)-HiBiT target engagement assay. DMSO-normalized luminescence of focused amine library reacted with **8** (mean ± s.e.m., *n* = 4) or vehicle (box and whiskers, min to max, *n* = 4). (e) Chemical structures of **9** and SR-0813 (**10**). (f) Activity of **9** and SR-0813 by HTRF (left) and ENL(YEATS)-HiBiT target engagement assay (right). Mean ± s.e.m., *n* = 4. (g) Binding curves for SR-0813 to ENL YEATS (left) and YEATS4 YEATS (right). RU, response units.

First, we appended iminosulfur oxydifluoride groups to the meta (**5**) and para (**6**) positions on the phenyl ring of **1** and confirmed that these compounds retained activity against ENL YEATS by HTRF (Figure S2a). We then reacted each compound in parallel with a library of 288 primary and secondary amines and screened the crude products at a putative concentration of 2 µM (Figure 2a,b). In general, meta substitutions seemed to be preferred over para, and while the parent molecules are inactive at 2 µM (Figure S2a), several crude reaction products fully inhibited ENL YEATS at this presumed dose (Figure 2b, Table S2). Selecting 28 crude products for dose-response examination, we observed striking improvements in potency compared to the parent molecule (Figure 2c, Table S3). The most potent hit – formed by the reaction of **5** with histamine – inhibited ENL YEATS with a half-maximal inhibitory concentration (IC_50_) of 120 nM by HTRF (Figure 2c). To confirm this result, the compound was resynthesized, purified, and then tested for activity against ENL YEATS (**7**, Figure S2b). By HTRF, the activity of the purified product matched closely with that of the crude (IC_50_ = 230 nM; Figure S2c). Compared to the initial screening hit (IC_50_ = 7 µM), this represents a 30-fold improvement in potency achieved in a single experiment. Nevertheless, this dramatic improvement in biochemical activity did not translate to similar gains in cell-based target engagement (EC_50_, **1** = 10 µM; EC_50_, **7** = 1.5 µM; Figure S2d).

Hoping to improve the performance of **7** in cells, we speculated that SuFEx-based screening is sufficiently biocompatible to permit testing of crude products directly in living cells. By this time, we had discovered the improvement in potency afforded by the cyclobutylamine modification in compound **3**, so we installed the iminosulfur oxydifluoride group onto the meta position of **3** instead of **1**, yielding compound **8** (Figure 2d). This SuFEx-compatible analog was then reacted in parallel with a small collection of amines sterically related to histamine and screened for target engagement in cells at 2 µM. A noticeable preference for piperidine analogs was revealed by this experiment, including one with the same CH_2_CH_2_ linker as the histamine in **7** (Figure 2d, Table S4). To compare the imidazole and piperidine analogs directly, we synthesized and purified compounds **9** and **10** (SR-0813) (Figure 2e). While **9** features superior biochemical potency (IC_50_, **9** = 6.9 nM; IC_50_, SR-0813 = 25 nM), SR-0813 engages the ENL YEATS domain more potently in cells and elicits a greater E_max_ (Figure 2f). Encouraged by these improvements, we next sought to evaluate the suitability of SR-0813 as a chemical probe for the ENL YEATS domain.

We first sought biophysical confirmation of ligand binding by surface plasmon resonance (SPR) and determined a 30 nM affinity between SR-0813 and the ENL YEATS domain (Figure 2g, Figure S2e). Then, we profiled SR-0813 for activity against other acyl-lysine readers, including bromodomains and other YEATS domains. This revealed potent activity against AF9 YEATS, the closest homolog of ENL,^12^ by both HTRF and CETSA (Figure S3a,b). In contrast, virtually no binding to the YEATS4 YEATS domain was observed by SPR (Figure 2g, Figure S3c,d), nor was any activity against BRD4 BD1 detected by HTRF (Figure S3e). In fact, not a single bromodomain off target was identified for SR-0813 using the BROMO*scan* profiling service from DiscoverX (Figure S4a; Table S5). Given the similarity of its imidazopyridine core to many bicyclic kinase inhibitors, we also tested SR-0813 for kinase off-targets using the KINOME*scan* profiling service (Figure S4b, Table S6). Of the 468 kinases tested, 4 were identified as potential off targets, but subsequent affinity determination revealed 3 of these as false-positive hits (Figure S4b,c). The remaining kinase, YSK4 (also known as MAP3K19), was bound by SR-0813 with low affinity (*K*_d_ = 3.5 µM) and would not be expected to complicate studies of ENL function in cellular model systems.

Armed with a highly potent and selective ENL YEATS inhibitor, we sought to benchmark its activity in comparison to past genetic experiments. Previously, alanine substitutions in the ENL YEATS domain helped to establish acyl-lysine recognition as an important contributor to ENL chromatin localization.^10,11^ To test this model via direct pharmacological occupation of the ENL YEATS domain, we preformed chromatin immunoprecipitation (ChIP) of ENL following treatment of MV4;11 cells with SR-0813 for 4 h. Quantification of ChIP enrichment by qPCR demonstrated clear dose-dependent eviction of ENL from known sites of ENL enrichment, including the *HOXA10* gene body and *MYB* promoter (Figure 3a). Downstream of these effects, we hypothesized that SR-0813 would suppress ENL-dependent transcriptional control. ENL has previously been shown to be important for *HOXA9/10, MEIS1*, and *MYC* gene expression, helping to enforce the differentiation block that is characteristic of MLL-rearranged leukemia.^10^ Kinetic investigation of these transcripts in response to SR-0813 treatment revealed rapid and prolonged suppression, beginning as early as 3 hours after drug exposure and sustained over the course of 3 days (Figure 3b). We also observed increased abundance of the *ITGAM* transcript, which encodes for CD11b (Figure 3b), consistent with previous RNA-seq studies that measured the effects of dTAG-mediated ENL degradation in MV4;11 cells.^10^ As noted above, CD11b is also elevated in MV4;11 upon SR-1114 treatment (Figure 1h) and in MOLM-13 cells upon genetic depletion of ENL by CRISPR/Cas9.^11^ Taken together, these data confirm the on-target activity of SR-0813 in living AML cells.

**Figure 3.**
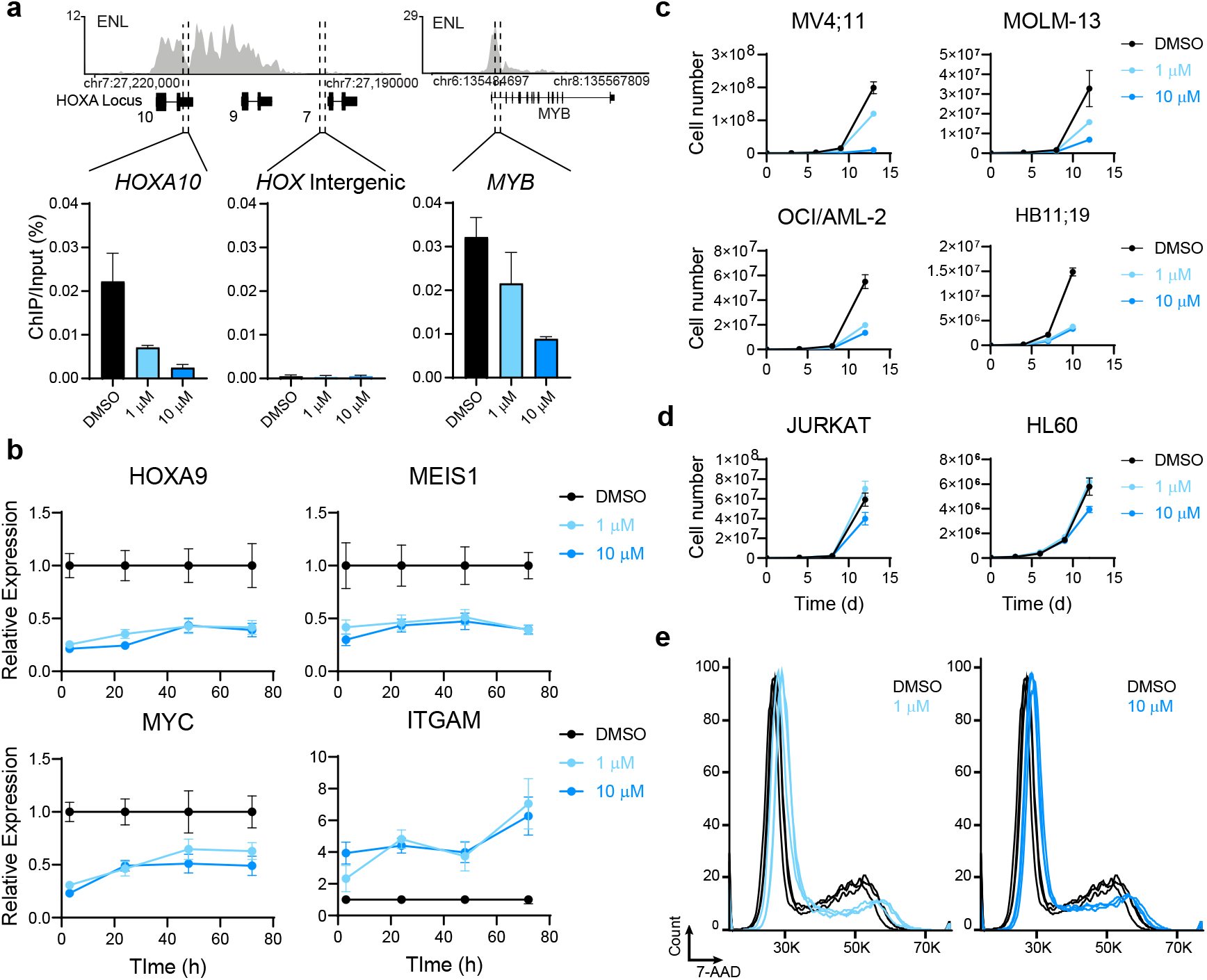
On-target effects of SR-0813 in leukemia. (a) ENL displacement from chromatin by SR-0813. Top: gene track views of ENL ChIP-seq signal from DMSO treated MV4;11cells. Bottom: ChIP-qPCR signal in response to SR-0813 treatment. Mean ± s.d. (b) Change in ENL target gene transcript abundance in MV4;11 cells treated with DMSO or SR-0813. qRT-PCR, *n* = 3. (c) Proliferation of MLL-fusion leukemia cell lines in response to SR-0813. Mean +-s.e.m. *n* = 3. (d) Proliferation of ENL-insensitive acute leukemia cell lines in response to SR-0813. (e) Cell cycle analysis of MV4;11 cells treated with DMSO or SR-0813 for 72 h. *n* = 3.

After confirming the most proximal biomarkers of ENL YEATS engagement in AML cells, we next characterized the effects of SR-0813 on acute leukemia proliferation, comparing its effects to the known pattern of sensitivity to ENL depletion by CRISPR/Cas9. We began by testing the response of 3 MLL-fusion leukemia that were previously shown to be sensitive to genetic loss of ENL – MV4;11 (MLL-AF4 AML), MOLM-13 (MLL-AF9 AML), and OCI/AML-2 (MLL-AF6 AML).^10,11^ In each of these three cell lines, SR-0813 elicited a dose-dependent inhibition of cellular proliferation (Figure 3c) that mirrored the dose-dependent displacement of ENL from chromatin (Figure 3a). Similarly, HB11;19 cells, which have not been tested for their reliance on wild-type ENL but harbor an MLL-ENL fusion, are also sensitive to SR-0813 exposure. In contrast, the growth patterns of JURKAT (T-ALL) and HL-60 (MLL wild-type AML) cells – both of which are insensitive to ENL loss^10^ – are unaffected by SR-0813 (Figure 3d). Moreover, we determined that the inhibition of MV4;11 growth by SR-0813 is accompanied by a decrease in the number of cells at the G2 stage of the cell cycle (Figure 3e), in line with previous data demonstrating G1 arrest upon loss of ENL.^10,11^ Taken together, these data are consistent with an on-target mechanism of growth inhibition resulting from the selective engagement of ENL YEATS.

To more fully assess the impact of SR-0813 on ENL-dependent gene control, we integrated genomic measures of ENL chromatin localization with dynamic gene expression profiling. ChIP followed by next generation DNA sequencing (ChIP-seq) revealed bulk displacement of ENL from chromatin upon treatment with SR-0813 (Figure 4a). The magnitude of ENL displacement genome-wide followed a similar dose-response pattern as was observed by ChIP-qPCR and in leukemia proliferation studies (Figure 4a). This is readily apparent both genome-wide and at specific sites that are relevant to AML pathogenesis, like the *HOXA* cluster, *MYC* promoter, and *MYC* enhancer (Figure 4a,b, Figure S5a). In previous work, we noted an asymmetric distribution of ENL on the genome whereby a small percentage of sites are occupied by a disproportionately large amount of ENL.^10^ These genes, in addition to being enriched with disease-relevant leukemia proto-oncogenes, are preferentially sensitive to acute loss of ENL by dTAG-mediated degradation. Here, we again detected this asymmetry in ENL localization (Figure 4c) and used 3’ mRNA sequencing to determine whether asymmetrically loaded ENL target genes are preferentially sensitive to SR-0813. After a 4-hour exposure in MV4;11 cells, SR-0813 elicited relatively few changes genome-wide, as expected (Figure 4d). In contrast, it produced a strikingly selective suppression of ENL target genes, with even more pronounced effects on those with disproportionate loads of ENL (Figure 4d,e). These genes include an impressive collection of transcription factors with well-established roles as leukemia drivers, including HOXA9/10, MEIS1, MYC, MYB, and ZEB2.^8,27–30^ Therefore, we show for the first time that direct pharmacological disruption of ENL chromatin localization can selectively suppress leukemogenic transcription.

**Figure 4.**
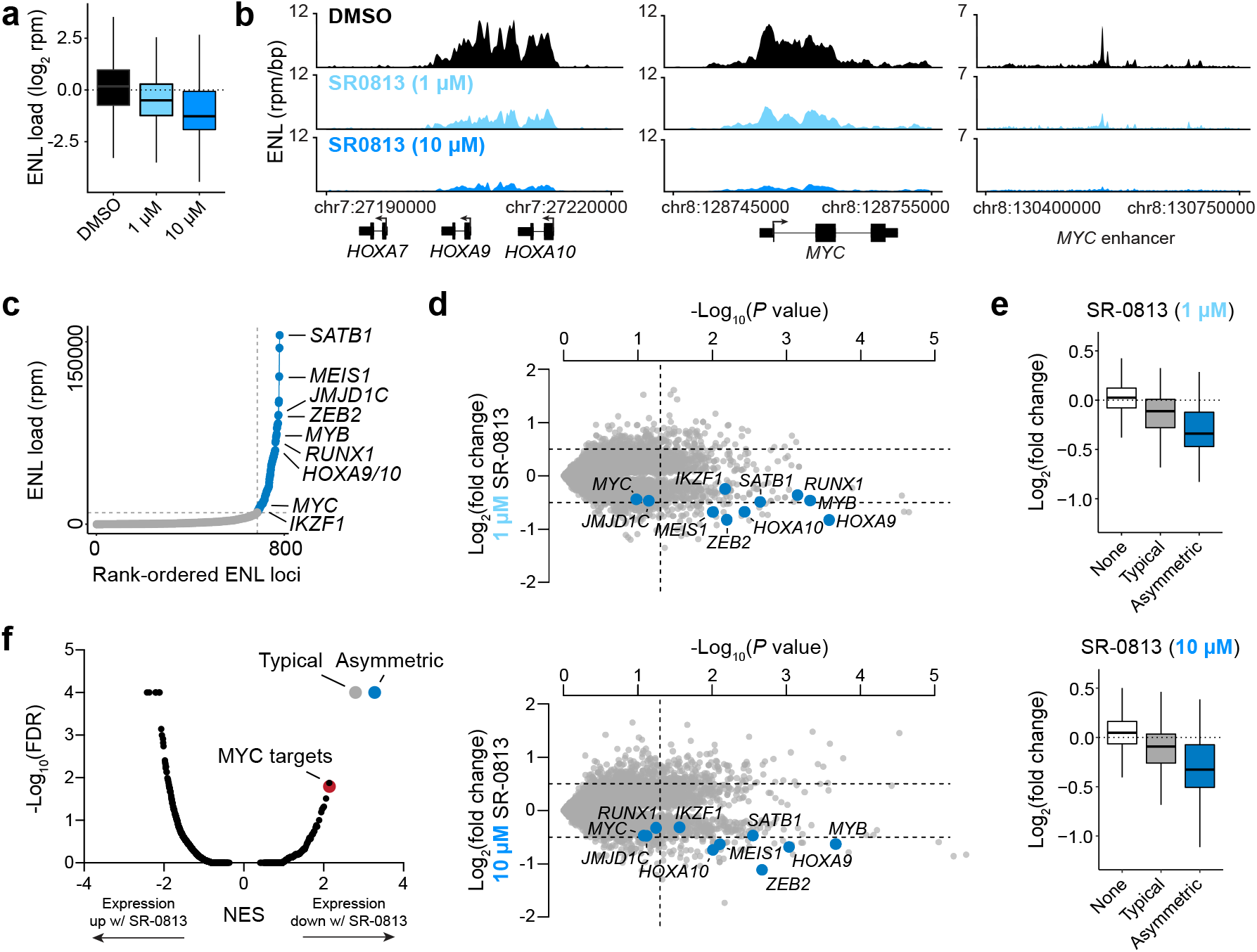
Selective suppression of ENL target genes by SR-0813. (a) Boxplots of ChIP-seq signal at ENL-bound loci (*n* = 2152) from MV4;11 cells treated with DMSO or SR-0813. (b) Gene tracks of ChIP-seq signal at asymmetrically loaded loci. (c) Rank-ordered plot of ENL ChIP-Seq signal at ENL peaks showing asymmetry in ENL binding to target genes. (d) Volcano plot of DMSO-normalized gene expression changes in MV4;11 cells treated with SR-0813 for 4 h (top: 1 µM, bottom: 10 µM). *P* value calculated with two-tailed Student’s t-test (*n* = 3). (e) Boxplots of gene expression changes at genes not bound by ENL (white, *n* = 12,163), typical ENL target genes (grey, *n* = 197), and asymmetrically loaded ENL target genes (blue, *n* = 74). (f) Gene set enrichment analysis (GSEA) of gene-expression changes with 4 h SR-0813 (10 µM) treatment.

To contextualize the selectivity of SR-0813-induced transcriptional effects for ENL target genes, we applied an unbiased gene set enrichment analysis (GSEA) to the 3’ mRNA-seq data set, profiling typical and asymmetric ENL targets along with 5,552 other gene sets in the molecular signatures database (MSigDB). Among these, typical and asymmetric ENL targets were by far the most highly enriched, followed next by a gene set related to *FLT3* internal tandem duplications (ITD)^31^ (a pathogenic alteration that is present in MV4;11 cells) and MYC targets (Figure 4f, Figure S5b). These unbiased data show that SR-0813-dependent changes in gene expression are exceptionally selective for ENL target genes. Collectively, we interpret these data as strong support for SR-0813 as an exquisitely selective chemical probe for the ENL YEATS domain.

## Discussion

In this study, we report on the discovery of SR-0813, a highly potent and selective chemical probe for ENL/AF9 YEATS domains. Our ligand discovery efforts were motivated by the previous identification of ENL as a critical dependency in acute leukemia.^10,11^ Using genetic tools to characterize this dependency *in vitro* and *in vivo*, we and others formulated the hypothesis that chromatin-competitive inhibition of the ENL YEATS domain would selectively inhibit the growth of ENL-dependent acute leukemia.^10,11^ While multiple reports of synthetic YEATS domain ligands have since followed,^16–21^ none have been used to study ENL-dependent transcriptional control or ENL-dependent leukemia proliferation. Here, we provide the first pharmacological validation of ENL as a leukemia target, reproducing key mechanistic insights made in the index target identification studies.

SR-0813 was elaborated from a high-throughput screening hit using an innovative high-throughput medicinal chemistry approach that leverages SuFEx reactivity to expedite chemical diversification. The conditions of these SuFEx transformations are highly biocompatible, enabling crude products to be tested directly in biochemical assays.^22^ Expanding on this approach, we now demonstrate that crude SuFEx products can also be tested directly in living cells, which should prove highly enabling for a number of cell-based drug discovery applications.

We were previously able to study the impact of ENL on leukemia gene control by using the dTAG system to elicit acute ENL degradation.^10^ These studies established ENL as a highly selective transcriptional co-activator that disproportionately affects the expression of several leukemia proto-oncogenes. Using the dTAG system in advance of embarking on a drug discovery campaign provided our group with a unique ability to anticipate the on-target effects of direct-acting ENL YEATS inhibitors. That is, we used the mechanistic insights from dTAG-based studies to benchmark the effects of our compounds on known aspects of ENL target biology. The success of this approach underscores the value of dTAG technology for target identification studies. Now equipped with a well-characterized chemical probe, we and others will be able to disrupt ENL function in a much broader range of systems than is possible with the dTAG system, which requires extensive biological engineering.

Here, we studied ENL-dependent gene control with direct-acting small molecules for the first time, integrating dynamic and unbiased measures of chromatin structure and function. In fact, this is the first study to demonstrate that small-molecules can displace endogenous ENL from chromatin, as previous studies judged target engagement with exogenous overexpression of tagged constructs and luminescence-based proximity reporters that require hyperacetylation of the genome via pan-HDAC inhibition.^17^ Since not all acetyl-lysine reader domains mediate chromatin localization of their host proteins (e.g. the bromodomains of SMARCA2/4 and CBP/p300),^32,33^ this provides an important insight into function of the ENL YEATS domain. Upon displacing ENL from chromatin, SR-0813 elicits a remarkably selective suppression of ENL target genes. Compared to over 5,000 other gene sets, ENL-bound genes are clearly the most severely affected by SR-0813 in MV4;11 cells. Moreover, the group of asymmetrically loaded ENL target genes, which were previously found to be hypersensitive to dTAG-mediated ENL degradation, were found to be identically sensitive to SR-0813 treatment. This represents the first evidence that direct-acting chemical tools can reproduce the genetic studies that established ENL as a selective transactivator of leukemia gene expression.

In replicating past genetic experiments, we simultaneously reinforce the evidence for ENL as a compelling leukemia dependency and validate SR-0813 as a useful chemical probe to study ENL/AF9 YEATS domain biology. We anticipate that SR-0813 and SR-1114 will represent timely additions to the collection of tools available for studying ENL and AF9 YEATS domains, as many aspects of ENL biology remain poorly understood. For instance, ENL stands in contrast to the activity of many other co-activators that are thought to be more global regulators of gene control. For instance, BRD4, an analogous acetyl-lysine reader and AML target is universally required for active gene transcription.^28,34,35^ While the selectivity of ENL-dependent gene control in AML tracks with its asymmetric localization on the genome, it is still not understood how that asymmetry is established. Our molecules will help to address these and other questions, which will broaden our understanding of the role that druggable co-regulators play in driving pathogenic gene control.

## Acknowledgement

We gratefully acknowledge L.L. Lairson for critical reading of the manuscript; G.E. Winter for helpful discussions regarding the work; and L.L. Lairson, P.G. Schultz, and I.A. Wilson for access to instrumentation. We also thank P.G. Schultz and the Schultz laboratory for helpful discussions and technical support. Sequencing was performed by the Scripps Research Next Generation Sequencing Core (La Jolla). This work was supported by the National Institutes of Health (NIH) through an NIH Director’s Early Independence Award (DP5-OD26380) to M.A.E and by the Leukemia and Lymphoma Society through a New Idea Award to M.A.E.

