## Supplemental Information for "Chemical inhibition of ENL/AF9 YEATS domains in acute leukemia"

**Affiliations:**

### Methods

**Protein production.** ENL and YEATS4 YEATS domains were recombinantly expressed and purified exactly as previously described in detail.[1] Briefly, ENL(1-148) was cloned into a modified pET21a (Novagen) expression vector with an N-terminal StrepII-SUMO or SUMO-AviTag and a C-terminal 6xHis tag and expressed in BL21(DE3) cells (New England Biolabs) by isopropyl-β-D-thiogalactopyranoside (IPTG) induction (0.4 mM) at 25 °C for 20 h. After purification by metal affinity chromatography (NiNTA agarose, Qiagen), buffer exchange to 100 mM Tris-HCl at pH 8.0, 150 mM NaCl, 1 mM EDTA, and 1 mM DTT using PD10 desalting columns (GE Life Sciences), and cleavage from SUMO by Ulp1 digestion, ENL YEATS(1-148) was recovered from the flow through of a StrepTrap column (GE Life Science). YEATS4(13-158) was cloned into pET28a with an N-terminal 6xHis and a C-terminal AviTag and expressed in BL21(DE3) cells by IPTG induction (0.25 mM) at 16 °C for 20 h. After purification by metal affinity chromatography (TALON Cobalt Resin, Takara) and buffer exchange to 50 mM sodium phosphate (pH 7.4) and 150 mM NaCl using Amnicon Ultracel 10K desalting columns (GE Life Sciences), YEATS4(13-158) was recovered by size exclusion chromatography using a HiLoad Superdex column (GE) on an AKTA pure (GE). To produce biotinylated protein, BL21(DE3) cells were co-transformed with the AviTag fusions and a biotin ligase expression vector, pBirAcm (Avidity). AF9 YEATS was purchased from ActiveMotif (Cat. No. 81124), and BRD4 BD1 was purchased from Cayman (Cat. No. 11720)

**Cell lines.** MV4;11 (RPMI 1640 supplemented with 10% Fetal Bovine Serum (FBS), penicillin and streptomycin), MOLM-13 (RPMI 1640 supplemented with 10% FBS, penicillin and streptomycin), OCI/AML-2 (RPMI 1640 supplemented with 10% FBS, penicillin and streptomycin), JURKAT (RPMI 1640 supplemented with 10% FBS, penicillin and streptomycin), HL-60 (RPMI 1640 supplemented with 20% FBS, penicillin and streptomycin) were provided by the laboratory of Prof. James E. Bradner. HB11;19 11 (RPMI 1640 supplemented with 10% Fetal Bovine Serum (FBS), penicillin and streptomycin) were provided by Prof. Akihiko Yokoyama. All cell lines were periodically (two to four times annually) tested for mycoplasma infections and never tested positive.

**HTRF.** AF9 YEATS, BRD4 BD1, and ENL YEATS HTRF assays were performed exactly as previously described.[1] All assays were conducted by combining recombinant AF9 YEATS (5 nM), BRD4 BD1 (10 nM), or ENL YEATS (5 nM), with synthetic histone peptide (100 nM $H3_{13\text{-}32}$K27cr custom synthesized by ABclonal, 13.3 nM tetra-acetylated H4 from BioVision Cat. No. 7144-01, or 13.1 nM $H3_{13\text{-}32}$K27cr, respectively), 1 nM LanthaScreen Eu-anti-His Tag antibody (ThermoFisher, PV5597) and 8.9 nM SureLight allophycocyanin-streptavidin (PerkinElmer, CR130-100). For the primary screen, the assay mixture was dispensed into black 1536-well plates (Greiner HiBase) at 6 μL/well and compounds were added by acoustic transfer (Labcyte, Echo Liquid Handler). Dose-response assays were performed with 10 or 20 μL assay volumes in black 384-well low-volume plates (Corning, Cat. No. 3821) and compounds were added by pin tool transfer (Biomek FX). Compounds were incubated with the assay mixture for at least 2 hours before measuring the HTRF signal on a PHERAstar plate reader (BMG Labtech; simultaneous dual emission; excitation = 337 nm, emission 1 = 665 nm, emission 2 = 620 nm). The HTRF signal was calculated as the ratio of emission 1 to emission 2. Each plate included samples treated with vehicle (DMSO) or lacking the peptide substrate, which were used to calculate the percent inhibition for compound-treated wells.

**Luminescence-based CETSA.** ENL(YEATS)-HiBiT and AF9(YEATS)-HiBiT constructs were stably expressed in OCI/AML-2 cells by lentiviral transduction and used for luminescence-based target engagement assays as previously described.[1] Briefly, cells were transferred to white 384-well tissue culture plates (Greiner Bio-One Cat. No. 781080) with 20,000 cells in 20 μL per well. Compounds were added by pin tool transfer (Biomek FX) and incubated for 2 hours before addition of 20 μL Nano-Glo HiBiT Lytic Reagent (Promega, HiBiT Lytic Detection System, Cat. No. N3040) per well. Luminescence was measured after 30 min after addition of the detection reagent using an EnVision multilabel plate reader (PerkinElmer, Model No. 2104).

**SPR.** Surface plasmon resonance experiments were performed on a Biacore S200 (Cytiva). A series S streptavidin sensor chip was used for ligand immobilization (Cytiva). Biotinylated ENL and biotinylated YEATS4 were prepared in HBS-EP+ buffer (10 mM HEPES, 150 mM NaCl, 3 mM EDTA, 0.05% P20, pH 7.4). Ligand immobilization was carried out in flow cell 2 and 4 with a series of ligand injections at a flow rate of 5 μL/min, until the desired density was achieved (1616 RU for ENL, and 2416 RU for YEATS4). Flow cells 1 and 3 were left blank as a reference. HBS-EP+ buffer with 5% DMSO was used as running buffer and for the preparation of all analyte samples. Analytes were serially diluted in polystyrene 96 well plates immediately prior to injection. Before analyte injection, 3 startup cycles were run with injections of running buffer over all flow cells. Binding data was collected by injecting the analyte at a flow rate of 30 μL/min over two flow cells (reference and immobilized ligand) at a temperature of 25 °C. The association to biotin-ENL or biotin-YEATS4 was measured over 60 seconds, and the dissociation over 180 seconds. The flow cell surfaces were regenerated with 30 second injections of running buffer over each flow cell. Duplicate injections were carried out for each sample, including the blank. Affinity curves, association and dissociation constants, and sensograms were determined using Biacore S200 evaluation software.

**Proteomics.** MV4;11 cells were treated with 10 μM SR-1114 for 4 h or 16 h and then flash frozen. Cell pellets were lysed in 150 μL of lysis buffer (10 mL of PBS with 1 tablet of Roche complete, mini, EDTA-free Protease Inhibitor Cocktail) using probe sonicator (3 x 10 pulses). Protein concentration was adjusted to 1.0 mg/mL. 100 μL of protein samples (100 μg protein) were transferred to new Eppendorf tubes (1.5 mL) containing urea (48 mg/tube, final concentration is 8 M). 5 μL of DTT (200 mM stock in water) was added to the tubes (final concentration is 10 mM). The samples were incubated at 65 °C for 15 min. After this, 5 μL of iodoacetamide (400 mM stock in water) was added to the tubes (final concentration is 20 mM). The samples were gently shake in the dark at 37 °C for 30 min. 600 μL of MeOH, 200 μL of $CHCl_3$, and 500 μL of water were added to the tubes. The mixture was vortexed and centrifuged (10,000 g, 10 min, 4 °C). A protein disc was formed at the interface of $CHCl_3$ and aqueous layers. After aspirating the top layer, 600 μL of MeOH was added and the proteins were pelleted (10,000 g, 10 min, 4 °C).

The protein pellets were resuspended in 160 μL of EPPS buffer (200 mM, pH 8). 4 μL of LysC solution (0.5 μg/μL in water) was added to each sample. The samples were incubated at 37 °C for 2 h. 10 μL of trypsin (0.5 μg/μL in trypsin buffer) and 1.8 μL of $CaCl_2$ (100 mM stock in water) were added. The samples were incubated at 37 °C with for 16 h.

Peptide concentration was determined. For each sample, a volume corresponding to 12.5 μg of peptides was transferred to a new Eppendorf tube and the total volume was brought up to 35 μL with EPPS buffer

(200 mM, pH 8). 9 μL of $CH_3CN$ was added to each sample. Then 3 μL of TMT tags (20 μg/μL in $CH_3CN$) was added and the samples were incubated at RT for 30 min. Additional TMT tag (3 μL/sample, 20 μg/μL) was added and the samples were incubated at RT for another 30 min. TMT labeling reaction was quenched by the addition of 6 μL of hydroxylamine (5% in water). After a 15 min incubation at RT, 2.5 μL of formic acid was added.

For ratio check, 2 μL of each sample were combined in a separate Eppendorf tube and dried using SpeedVac vacuum concentrator. The residue was dissolved in 20 μL of buffer A (5% $CH_3CN$ in water with 0.1% formic acid) and desalted using C18 stage tips. The residue was dissolved in 10 μL of buffer A and analyzed by mass-spectrometry using the following LC-MS gradient: 5% buffer B (95% $CH_3CN$ in water with 0.1% formic acid) in buffer A from 0-15 min, 5-15% buffer B from 15-17.5 min, 15-35% buffer B from 17.5-92.5 min, 35-95% buffer B from 92.5-95 min, 95% buffer B from 95-105 min, 95-5% buffer B from 105-107 min, and 5% buffer B from 107-125 min and standard MS3-based quantification described below. Ratios were determined from the average peak intensities corresponding to each channel. Samples (20 μL/sample, final volumes adjusted based on the determined ratios) were combined in a new Eppendorf tube and dried using SpeedVac. The residue was dissolved in 500 μL of buffer A, desalted by Sep-Pak C18 cartridges, and subjected to HPLC fractionation.

The sample was fractionated into a 96 deep-well plate using a capillary column (ZORBAX 300Extend-C18, 3.5 μm) and separated at a flow rate of 0.5 mL/min using the following gradient: 100% buffer A from 0-2 min, 0-13% buffer C (10 mM aqueous $NH_4HCO_3$) from 2-3 min, 13-42% buffer C from 3-60 min, 42-100% buffer C from 60-61 min, 100% buffer C from 61-65 min, 100-0% buffer C from 65-66 min, 100% buffer A from 66-75 min, 0-13% buffer C from 75-78 min, 13-80% buffer C from 78-80 min, 80% buffer C from 80-85 min, 100% buffer A from 86-91 min, 0-13% buffer C from 91-94 min, 13-80% buffer C from 94-96 min, 80% buffer C from 96-101 min, and 80-0% buffer C from 101-102 min. Every 12th fraction was combined. The solvent was removed using SpeedVac vacuum concentrator. The resulting 12 fractions were dissolved in 15 μL of buffer A.

Samples were analyzed by liquid chromatography tandem mass-spectrometry using an Orbitrap Fusion mass spectrometer (Thermo Scientific) coupled to an UltiMate 3000 Series Rapid Separation LC system and autosampler (Thermo Scientific Dionex). The peptides were eluted onto a capillary column (75 μm inner diameter fused silica, packed with C18 (Waters, Acquity BEH C18, 1.7 μm, 25 cm) and separated at a flow rate of 0.25 μL/min using the following gradient: 5% buffer B in buffer A from 0-15 min, 5-35% buffer B from 15-155 min, 35-95% buffer B from 155-160 min, 95% buffer B from 160-169 min, 95-5% buffer B from 169-170 min, and 5% buffer B from 170-200 min. The voltage applied to the nano-LC electrospray ionization source was 1.9 kV. The scan sequence began with an MS1 master scan (Orbitrap analysis, resolution 120,000, 400-1700 m/z, RF lens 60%, automatic gain control [AGC] target 2E5, maximum injection time 50 ms, centroid mode) with dynamic exclusion enabled (repeat count 1, duration 15s). The top ten precursors were then selected for MS2/MS3 analysis. MS2 analysis consisted of: quadrupole isolation (isolation window 0.7) of precursor ion followed by collision-induced dissociation (CID) in the ion trap (AGC 1.8E4, normalized collision energy 35%, maximum injection time 120 ms). Following the acquisition of each MS2 spectrum, synchronous precursor selection (SPS) enabled the selection of up to 10 MS2 fragment ions for MS3 analysis. MS3 precursors were fragmented by HCD and analyzed using the Orbitrap (collision energy 55%, AGC 1.5E5, maximum injection time 120 ms, resolution was 50,000). For MS3 analysis, charge state-dependent isolation windows were used. For charge state z = 2, the MS isolation window was set at 1.2; for z = 3-6, the MS isolation window was set at 0.7.

RAW data was searched in Integrated Proteomics Pipeline (IP2) using the ProLuCID algorithm (publicly available at http://fields.scripps.edu/downloads.php) and a reverse concatenated, non-redundant variant of the Human UniProt database (release-2012_11). Cysteine residues were searched with a static modification for carboxyamidomethylation (+57.02146). Lysine residues and peptide N-termini were searched with a static modification for TMT-tag labeling (+229.162932). Methionine residues were searched with a dynamic modification for oxidation (+15.9949). MS3 quantification was performed using 10-plex TMT analysis parameters (m/z 126.127726, 127.124761, 127.131081, 128.128116, 128.134436, 129.131471, 129.13779, 130.134825, 130.141145 and 131.13818) with the mass tolerance of 20 ppm. Protein relative abundance was calculated using the corresponding MS3 intensity.

**Immunoblotting.** Whole cell lysates were prepared by solubilization of cell pellets in CelLytic M (Sigma Cat# C2978) 1X HALT protease inhibitor cocktail (Thermo) and 0.1% Benzonase (EMD Millipore) by vertexing 20s followed by 15 minute incubation on ice. Cleared lysates were prepared by centrifugation 18,000 g 11 minutes at 4° C. All samples were run with the same amount of total protein as determined by a BCA protein assay kit (Pierce). ENL (Cell Signaling Technology, #14893S) and GAPDH (Santa Cruz Biotechnology, sc-25778) antibodies were used at 1:1000 into 5% non-fat milk in 1X TBST and detected with fluorescently labeled infrared secondary antibodies (IRDye) on the Odyssey CLx Imager (LI-COR). Quantitation preformed in ImageStudio Lite (LI-COR) and nonlinear fit for $DC_{50}$ determination GraphPad Prism version 8.4.3 for Windows (GraphPad Software).

**DiscoverX profiling services.** Bromodomain and kinase domain off-target effects of SR-0813 (10 μM) were profiled by the commercially available BROMO*scan* (using the bromoMAX panel of 32 targets) and KINOME*scan* (using the scanMAX panel of 468 targets) profiling serves from DiscoverX.

**Cell proliferation and cell cycle assays.** Cell proliferation assays were carried out in 96-well tissue culture plates at 20,000 cells/well for all lines except for HL60s (Which were started at 40,000 cells/well) in 200μL with compounds added as 1:1000 dilutions of DMSO stocks, in triplicate. Culture density determined every 3-4 days using the Countess automated cell counter (Invitrogen), after which 20,000 live cells are reseeded in fresh media and compound. Cumulative cell count achieved by back calculation. For cell cycle analysis 500k cells were cultured in 1mL/well in a 24-well plate treated 1:1000 with compound stocks in DMSO or 1:1000 DMSO, in triplicate. 7-AAD staining was performed using the FITC BrdU Flow Kit from BD Biosciences (cat#). 7-AAD signal was analyzed by flow cytometry with a NovoCyte 3000 from Agilent. Flow cytometry results analyzed by FlowJo software (version 10.7 Becton, Dickinson and Company).

**qRT-PCR.** RNA extracts were prepared with the RNeasy Mini plus kit (Qiagen). cDNA was synthesized with the SuperScript VILO cDNA Synthesis Kit (Life Technologies cat# 11755050) at 100ng/uL RNA. cDNAs were analyzed in triplicate with a 7900HT Fast Real-Time PCR System (Applied Biosystems) in SYBR Select Master Mix (Life Technologies Cat# 4472908 ) with the primer pairs detailed below. Cycle numbers (Ct) determined by default setting on Design and Analysis (Applied Biosystems) and converted to expression fold change by normalization to B2M transcript levels.

**Primer Pairs for qRT-PCR.** HOXA9: forward 5'-GTATAGGGGCACCGCTTTTT-3', reverse 5'-AATGCTGAGAATGAGAGCGG-3'. MEIS1: forward 5'-CACGCTTTTTGTGACGCTT-3' reverse 5'-GGACAACAGCAGTGAGCAAG'3'. MYC: forward 5'-CACCGAGTCGTAGTCGAGGT-3' reverse 5'-TTTCGGGTAGTGGAAAACCA-3'. ITGAM: forward 5'-CAAAATACTGGAGCCTGGGA-3' reverse 5'-CCTGTTTCACGGAACCTCAG-3'. B2M: forward 5'- AATGTCGGATGGATGAAACC-3' reverse 5'-TAGCTGTGCTCGCGCTACT-3'.

**ChIP.** ChIP-seq was performed as previously described.[2] For ENL ChIPs, 100 million cells treated per condition in 100mL. Cells crosslinked for 10minutes rocking at RT by addition of 1/10$^{th}$ volume 10X crosslinking solution (11% formaldehyde, 50 mM HEPES pH 7.3, 100 mM NaCl, 1 mM EDTA pH 8.0, 0.5 mM EGTA pH 8.0). At the end of 10 minutes cultures were quenched in 125mM Glycine, freshly made, and rocked for an additional 10 minutes. Crosslinked cells were then washed 3X in cold 1X DPBS pH7.4 and pellets flash frozen in liquid nitrogen and stored at at -80° C. Pellets resuspended in cold lysis buffer 1 (LB1; 5 mL per 50 million cells; 50 mM HEPES pH 7.3, 140 mM NaCl, 1 mM EDTA, 10% glycerol, 0.5% NP- 40, and 0.25% Triton X-100, 1X HALT protease inhibitor cocktail from Thermo), and rotated end over end for 10 minutes at 4° C. Pellet collected at 1350 g and resuspended in cold lysis buffer 2 (LB2; 5 mL per 50 million cells; 10 mM Tris-HCl pH 8.0, 200 mM NaCl, 1 mM EDTA pH 8.0 and 0.5 mM EGTA pH 8.0, 1X HALT protease inhibitor cocktail from Thermo), and rotated end over end 10 minutes at 4° C.. Pellets collected at 1350 g and resuspended in 3mL sonication buffer (50 mM HEPES pH 7.3, 140 mM NaCl, 1 mM EDTA, 1 mM EGTA, 1% Triton X-100, 0.1% Na-deoxycholate, 0.5% SDS, 1X HALT protease inhibitor cocktail from Thermo), and split into 250uL volumes in 1.5mL Bioruptor Plus TPX microtubes (Diagenode, #C30010010) and sheared at 4° C using a waterbath sonicator (Bioruptor Pico, Diagenode; 25 minutes; 30 seconds on, 30 seconds off). Sheared lysates cleared by centrifugation at 18,000 g 11 minutes at 4° C and pooled. 50μL of the cleared lysate stored at -20° C for isolation as the input. The remaining cleared lysate was diluted 1:5 in cold sonication buffer lacking SDS, such that the final SDS concentration is 0.1%. Magnetic protein G beads (Dynabeads, ThermoFisher Scientific) were washed 3 times in 1mL of cold blocking buffer (1X DPBS +5mg/mL BSA), then resuspended in 1mL of blocking buffer with 10ug anti-ENL (Cell Signaling Technology, #14893S) and rotated end over end at 4° C overnight. Antibody-bead complexes washed 3x in cold blocking buffer and added to the diluted cleared sonicated lysates and rotated end over end overnight at 4° C. Bead bound fraction washed twice in cold sonication buffer (no SDS), once with 1 mL cold sonication buffer supplemented with 500 mM NaCl, once with cold LiCl wash buffer (20 mM Tris pH 8.0, 1 mM EDTA, 250 mM LiCl, 0.5% NP-40, 0.5% Na-deoxycholate), and once with TE supplemented with 50 mM NaCl. The remaining bound fraction was eluted by addition of 210μL elution buffer (50 mM Tris-HCl pH 8, 10 mM EDTA, and 1% SDS) and vortexing every 5 minutes while otherwise incubating at 65° C for a total of 15 min. Beads were then centrifuged at 18,000 g 1 minute and the supernatant removed and incubated overnight alongside the 50μL input sample after addition of 150μL elution buffer to bring up to 200μL for all samples. RNA was digested with 0.2 mg/mL RNase A (Roche, 10109169001) at 37° C for 2 hours and protein was digested with 0.2 mg/mL proteinase K (Life Technologies, AM2546) at 55° C for 30 minutes. DNA was isolated with phenol chloroform extraction and ethanol precipitation. Libraries for Illumina sequencing were prepared using ThruPLEX DNA-seq Kit (Takara cat#R400675) using 0.75ng of DNA and amplifying according to manufacturer instructions using the DNA Single Index Kit 12 Set A (Takara cat#R400695). Amplified libraries were isolated using AMPure beads (Agencourt AMPure XP). Library quantity and size distribution determined by Qubit™ 1X dsDNA HS Assay Kit (ThermoFisher cat#Q33231) and D1000 high sensitivity DNA tape run on a 4150 TapeStation System (Agilent); samples were multiplexed with equimolar DNA content, and sequenced on an Illumina NextSeq 500 (single end 75 bp reads).

**ChIP-qPCR.** ChIP-seq DNAs from the above method were diluted and subjected to qPCR interrogation. DNAs were analyzed in triplicate with a 7900HT Fast Real-Time PCR System (Applied Biosystems) in SYBR Select Master Mix (Life Technologies Cat# 4472908 ) with the primer pairs detailed below. Cycle numbers (Ct) determined by default setting on Design and Analysis (Applied Biosystems) and converted to expression fold change

**ChIP-qPCR Primers**. HOXA10: forward 5’-GCCGCTCTCGAGTAAGGTAC-3', reverse 5’-GGCAAAGAGTGGTCGGAAGA-3'. HOXA_intergenic_1: forward 5’-CGCCTTCCAGAGATTTGGGT-3', reverse 5’-TGTCTTTGCACCCAGGCTAG-3'. MEIS1_1: forward 5’-GGGAAACGCGAGCTTTTGTT-3', reverse 5’-GGCTAGGAAGTCCGCAAAGT-3'. MYB_1: forward 5’-TGCAAAGTTCAAAGGCGAGC-3', reverse 5’-CTTCTGCAGGACCTGAAGGG-3'.

**Quant-seq.** For Quantitative RNA-seq, cells were treated in triplicate with 1:1000 DMSO, 10uM SR-0813 or 1uM SR-0813 (from dilution of 1000X stocks in DMSO) treated for 4 hours and harvested. 500k cells resuspended in Lysis buffer containing ERCC spike in (SIRV set 3, Lexogen cat# K05103-0-0100), samples isolated with RNeasy Plus kit from Qiagen. 400ng of RNA used to prepare libraries for sequencing with QuantSeq 3‘ mRNA-Seq Library Prep Kit (FWD) for Illumina (Lexogen, cat#015.2) and equimolar multiplexed libraries were sequenced on an Illumina NextSeq 500 (single end 75 bp reads).

**ChIP-seq analysis.** ChIP datasets were aligned to human genome build hg19 with Bowtie2 (Version 2.2.9). All default settings were used except for -N 1 (allowing 1 mismatch in seed alignments). SAM alignment files were converted to BAM files with Samtools (Version 1.9) view. BAM files were then sorted and indexed with Samtools. Model-based Analysis of ChIP-seq (MACS) peak-finding algorithm (Version 1.4.2) was used to identify regions of ChIP-seq signal that were significantly enriched over background. Normalized read density for specific genomic loci of interest was determined using Bamliquidator (http://github.com/BradnerLab/pipeline/wiki/bamliquidator) read density calculator. ROSE2 was used to identify regions of asymmetric ENL binding (https://github.com/linlabbcm/rose2).

**Quant-seq analysis.** For Quant-seq analysis, quality of sequencing data was assured with Fastqc. SlamDunk (with the -q option to ignore SLAM-seq scoring) was used to align reads to human genome build hg19 and calculate counts per million (CPM) at all 3' UTRs. Average log2-transformed fold change values were calculated from the means of triplicate treatments. Transcripts with an average CPM < 3 were excluded from analysis.

**Synthetic procedures.**

**General**

All reagents and solvents were purchased from commercial suppliers and were used without further purification. $^{1}H$, $^{13}C$, and $^{19}F$ NMR spectra were collected using a Bruker 600, 500, or 400 MHz spectrometer with chemical shifts reported relative to residual deuterated solvent peaks or a tetramethylsilane internal standard. $CFCl_3$ was used as an internal standard for $^{19}F$-NMR. Accurate masses were measured using an ESI-TOF (HRMS, Agilent MSD) or MSQ Plus mass spectrometer (LRMS, Thermo Scientific). Reactions were monitored on TLC plates (silica gel 60, F254 coating, EMD Millipore, 1057150001), and spots were either monitored under UV light (254 mm) or stained with phosphomolybdic acid. The purity of the compounds that were tested in the biological assay was >95% based on $^{1}H$ NMR and reverse phase HPLC-UV on monitoring absorption at 240 nm.

**Representative procedure for the synthesis of imidazopyridine**

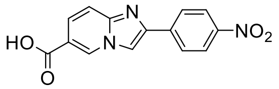


Exact Mass: 283.06
Molecular Weight: 283.24

To a DMF solution of 6-aminonicotinic acid (1 g, 7.24 mmol) was added 2-bromo-4-nitroacetophenone (3.5 g, 14.5 mmol, 2 eq.) and stirred overnight at 60°C. The reaction mixture was cooled to RT and to this solution was added ethyl acetate and water. The resulting precipitate was collected by filtration then washed with water and ethyl acetate to give a fairly pure target molecule as a red powder. The compound was used into the next step without further purification (2-(4-nitrophenyl)imidazo[1,2-*a*]pyridine-6-carboxylic acid, 1.39 g, 4.9 mmol, 68%). LRMS (+) calcd for $(M+H)^+$ 284.1. Found 284.2. $^{1}H$ NMR (600 MHz, DMSO-$d_6$) δ 9.27 (t, *J* = 1.4 Hz, 1H), 8.76 (s, 1H), 8.36 – 8.32 (m, 2H), 8.23 (dd, *J* = 9.1, 2.4 Hz, 2H), 7.73 – 7.66 (m, 2H). $^{13}C$ NMR (151 MHz, DMSO-$d_6$) δ 165.8, 146.8, 145.6, 143.3, 139.7, 131.3, 126.6, 125.3, 124.3, 117.1, 116.3, 112.9.

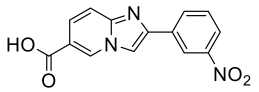


2-(3-nitrophenyl)imidazo[1,2-*a*]pyridine-6-carboxylic acid, 1.43 g (5.1 mmol, 70%). LRMS (+) calcd for $(M+H)^+$ 284.1. Found 284.2. $^{1}H$ NMR (600 MHz, DMSO-$d_6$) δ 9.23 (t, *J* = 1.4 Hz, 1H), 8.76 (t, *J* = 2.0 Hz, 1H), 8.74 (s, 1H), 8.39 (dt, *J* = 7.7, 1.3 Hz, 1H), 8.20 (ddd, *J* = 8.2, 2.4, 1.0 Hz, 1H), 7.79 – 7.76 (m, 1H), 7.70 – 7.68 (m, 2H). $^{13}C$ NMR (151 MHz, DMSO-$d_6$) δ 165.8, 148.4, 145.4, 143.3, 135.0, 131.9, 131.2, 130.5, 125.1, 122.6, 122.4, 119.9, 116.2, 111.8.

**Representative procedure for the amide coupling**

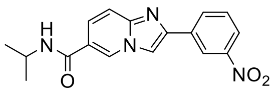


Exact Mass: 324.12
Molecular Weight: 324.34

To a DMF solution of 2-(3-nitrophenyl)imidazo[1,2-a]pyridine-6-carboxylic acid (1.5 g, 5.3 mmol) was added *N,N*-diisopropylethylamine (2.8 mL, 2 g, 15.5 mmol) followed by 1-[Bis(dimethylamino)methylene]-1H-1,2,3-triazolo[4,5-*b*]pyridinium 3-oxid hexafluorophosphate (2.4 g, 6.32 mmol, 1.2 eq.) and isopropylamine (470 mg, 8 mmol, 1.5 eq.) and stirred overnight at RT. Solvent was removed and to the residue was added ethyl acetate. The solid was collected by filtration and washed with ethyl acetate to give a fairly pure target molecule *N*-isopropyl-2-(3-nitrophenyl)imidazo[1,2-*a*]pyridine-6-carboxamide as a yellow powder (860 mg, 2.65 mmol, 50%). LRMS (+) calcd for $(M+H)^+$ 325.1. Found 325.2. $^{1}H$ NMR (600 MHz, DMSO-$d_6$) δ 9.11 – 9.04 (m, 1H), 8.78 (t, *J* = 2.0 Hz, 1H), 8.72 (s, 1H), 8.45 – 8.36 (m, 2H), 8.19 (d, *J* = 8.4 Hz, 1H), 7.80 – 7.71 (m, 2H), 7.67 (d, *J* = 9.6 Hz, 1H), 4.20

– 4.09 (m, 1H), 1.21 (d, *J* = 6.6 Hz, 6H). $^{13}$C NMR (151 MHz, DMSO-$d_6$) δ 163.0, 148.4, 145.1, 143.2, 135.3, 131.9, 130.4, 128.2, 124.3, 122.5, 120.6, 119.8, 115.9, 111.5, 41.2, 22.3.

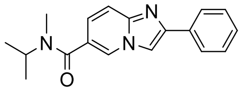

Exact Mass: 293.15
Molecular Weight: 293.37

*N*-isopropyl-*N*-methyl-2-phenylimidazo[1,2-*a*]pyridine-6-carboxamide. LRMS (+) calcd for (M+H)$^+$ 294.2. Found 294.1. $^1$H NMR (400 MHz, DMSO-$d_6$) δ 8.70 (s, 1H), 8.43 (s, 1H), 7.98 (d, *J* = 7.5 Hz, 2H), 7.63 (d, *J* = 9.2 Hz, 1H), 7.46 (t, *J* = 7.6 Hz, 2H), 7.35 (t, *J* = 7.3 Hz, 1H), 7.26 (t, *J* = 8.9 Hz, 1H), 4.70 – 4.10 (broad m, 1H), 2.86 (s, 3H), 1.18 (d, *J* = 8.0 Hz, 6H).

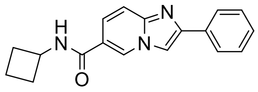

Exact Mass: 291.14
Molecular Weight: 291.35

*N*-cyclobutyl-2-phenylimidazo[1,2-*a*]pyridine-6-carboxamide. LRMS (+) calcd for (M+H)$^+$ 292.1. Found 292.1. $^1$H NMR (400 MHz, DMSO-$d_6$) δ 9.06 (s, 1H), 8.73 (d, *J* = 7.4 Hz, 1H), 8.50 (s, 1H), 7.98 (d, *J* = 7.3 Hz, 2H), 7.69 – 7.60 (m, 2H), 7.46 (t, *J* = 7.6 Hz, 2H), 7.35 (t, *J* = 7.3 Hz, 1H), 4.47 – 4.41 (m, 1H), 2.27 – 2.21 (m, 2H), 2.11 – 2.05 (m, 2H), 1.73 – 1.65 (m, 2H).

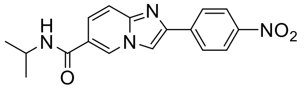

Exact Mass: 324.12
Molecular Weight: 324.34

*N*-isopropyl-2-(4-nitrophenyl)imidazo[1,2-*a*]pyridine-6-carboxamide. LRMS (+) calcd for (M+H)$^+$ 325.1. Found 325.2. $^1$H NMR (600 MHz, DMSO-$d_6$) δ 9.09 (d, *J* = 1.7 Hz, 1H), 8.73 (s, 1H), 8.42 – 8.21 (m, 5H), 7.80 – 7.61 (m, 2H), 4.18 – 4.06 (m, 1H), 1.20 (d, *J* = 6.6 Hz, 6H). $^{13}$C NMR (151 MHz, DMSO-$d_6$) δ 163.0, 146.7, 145.3, 143.2, 140.1, 128.3, 126.5, 124.6, 124.2, 120.7, 116.0, 112.6, 41.2, 22.3.

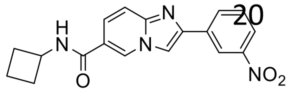

Exact Mass: 336.12
Molecular Weight: 336.35

*N*-cyclobutyl-2-(3-nitrophenyl)imidazo[1,2-*a*]pyridine-6-carboxamide (914 mg, 2.72 mmol, 51%). LRMS (+) calcd for (M+H)$^+$ 337.1. Found 337.2. $^1$H NMR (600 MHz, DMSO-$d_6$) δ 9.08 (q, *J* = 1.4 Hz, 1H), 8.82 – 8.75 (m, 2H), 8.73 (d, *J* = 1.3 Hz, 1H), 8.42 (dt, *J* = 7.8, 1.4 Hz, 1H), 8.19 (ddt, *J* = 8.2, 2.5, 1.2 Hz, 1H), 7.80 – 7.70 (m, 2H), 7.68 (d, *J* = 9.4 Hz, 1H), 4.50 – 4.40 (m, 1H), 2.28 – 2.22 (m, 2H), 2.14 – 2.11 (m, 2H), 1.78 – 1.63 (m, 2H). $^{13}$C NMR (151 MHz, DMSO-$d_6$) δ 162.9, 148.4, 145.1, 143.2, 135.3, 131.8, 130.4, 128.3, 124.3, 122.5, 120.3, 119.8, 115.9, 111.5, 44.6, 30.1, 14.8.

**Representative procedure for reduction of nitrophenyl into aniline**

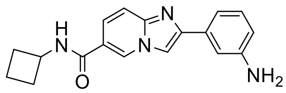

Exact Mass: 306.15
Molecular Weight: 306.37

To a DMF solution of *N*-cyclobutyl-2-(3-nitrophenyl)imidazo[1,2-*a*]pyridine-6-carboxamide (300 mg, 0.89 mmol) was added 5% Pd/C (100 mg), purged with $H_2$ gas and stirred at RT overnight. Solvent was removed and to the residue was added a solution of water:ACN:TFA = 90:10:0.1 and filtered through 0.22 μm PTFE syringe filter. The filtrate was purified by HPLC to give target molecule 2-(3-aminophenyl)-*N*-cyclobutylimidazo[1,2-*a*]pyridine-6-carboxamide as a white solid (135 mg, 0.44 mmol, 50%). LRMS (+) calcd for $(M+H)^+$ 307.2. Found 307.3. $^1$H NMR (600 MHz, DMSO-$d_6$) δ 9.21 (s, 1H), 8.91 (d, *J* = 7.4 Hz, 1H), 8.59 (s, 1H), 7.97 (d, *J* = 9.5 Hz, 1H), 7.78 (d, *J* = 9.4 Hz, 1H), 7.46 (d, *J* = 12.0 Hz, 2H), 7.37 (t, *J* = 7.8 Hz, 1H), 6.96 (d, *J* = 7.9 Hz, 1H), 4.45 (h, *J* = 8.1 Hz, 1H), 2.34 – 2.20 (m, 2H), 2.20 – 1.99 (m, 2H), 1.71 (ddd, *J* = 15.6, 10.3, 7.4 Hz, 2H). $^{13}$C NMR (151 MHz, DMSO-$d_6$) δ 162.3, 142.9, 140.9, 131.0, 130.2, 129.1, 127.6, 122.2, 119.1, 117.0, 115.4, 115.1, 113.7, 111.1, 44.8, 30.0, 14.8.

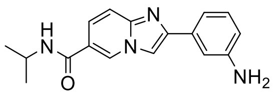


Exact Mass: 294.15
Molecular Weight: 294.36

2-(3-aminophenyl)-*N*-isopropylimidazo[1,2-*a*]pyridine-6-carboxamide. LRMS (+) calcd for $(M+H)^+$ 295.2. Found 295.3. $^1$H NMR (500 MHz, DMSO-$d_6$) δ 9.04 (dd, *J* = 1.8, 1.0 Hz, 1H), 8.35 – 8.30 (m, 2H), 7.65 (dd, *J* = 9.4, 1.8 Hz, 1H), 7.56 (d, *J* = 9.4 Hz, 1H), 7.24 (t, *J* = 1.9 Hz, 1H), 7.13 – 7.04 (m, 2H), 6.54 (dt, *J* = 7.2, 2.0 Hz, 1H), 5.15 (s, 2H), 4.12 (dq, *J* = 13.6, 6.6 Hz, 1H), 1.19 (d, *J* = 6.7 Hz, 6H). $^{13}$C NMR (151 MHz, DMSO-$d_6$) δ 163.3, 148.9, 146.4, 144.7, 134.0, 129.1, 127.9, 123.3, 120.0, 115.4, 113.8, 113.6, 111.2, 109.5, 41.2, 22.3.

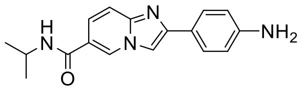


Exact Mass: 294.15
Molecular Weight: 294.36

2-(4-aminophenyl)-*N*-isopropylimidazo[1,2-*a*]pyridine-6-carboxamide. LRMS (+) calcd for $(M+H)^+$ 295.2. Found 295.3. $^1$H NMR (600 MHz, DMSO-$d_6$) δ 8.98 (d, *J* = 1.7 Hz, 1H), 8.31 (d, *J* = 7.6 Hz, 1H), 8.21 (s, 1H), 7.67 – 7.62 (m, 2H), 7.61 (dd, *J* = 9.4, 1.8 Hz, 1H), 7.51 (d, *J* = 9.3 Hz, 1H), 6.64 – 6.58 (m, 2H), 5.28 (s, 2H), 4.11 (dq, *J* = 13.5, 6.7 Hz, 1H), 1.18 (d, *J* = 6.6 Hz, 6H). $^{13}$C NMR (151 MHz, DMSO-$d_6$) δ 163.4, 148.9, 147.0, 144.7, 127.5, 126.8, 122.9, 121.16, 119.6, 114.9, 113.8, 107.5, 41.1, 22.4.

**Representative procedure for $SOF_4$ reaction**

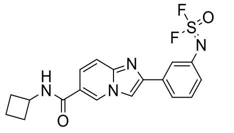


Exact Mass: 390.10
Molecular Weight: 390.41

A 25 mL round bottom flask equipped with a magnetic stir bar was charged with aniline (30.6 mg, 0.100 mmol), diisopropylethyl amine (DIPEA, 38.8 mg, 52 μL, 0.300 mmol, 3.00 equiv). Acetonitrile (MeCN, 1.0 mL) and *N*,*N*-dimethylformamide (DMF, 1.0 mL) were added. Topped by a rubber septum, the flask was evacuated (solvent gently bubbling) and backfilled with thionyl tetrafluoride ($O{=}SF_4$). The mixture was stirred vigorously at room temperature for 30 min. The solution was then purified by reversed phase preparative high-performance liquid chromatography (HPLC) using a water/MeCN containing 0.05% Vol.% trifluoroacetic acid (TFA) elution. The target compound (3-(6-(Cyclobutylcarbamoyl)imidazo[1,2-*a*]pyridin-2-yl)phenyl)sulfurimidoyl difluoride was obtained as an off-white solid (TFA salt, 10.1 mg, 0.0200 mmol, 20%). LRMS (+) calcd for $(M+H)^+$ 390.1, found 390.1. $^1$H NMR (600 MHz, DMSO-$d_6$) *δ* 9.22 (s, 1H), 8.95 (d, *J* = 7.4 Hz, 1H), 8.77 (s, 1H), 8.04 (dd, *J* = 9.4, 1.7 Hz, 1H), 7.91 (d, *J* = 9.4 Hz, 1H), 7.89 – 7.85 (m, 2H), 7.62 (t, *J* = 7.8 Hz, 1H), 7.35 (dd, *J* = 7.8, 2.0 Hz, 1H), 4.45 (q, *J* = 7.8 Hz, 1H), 2.30 – 2.21 (m, 2H), 2.10 (pd, *J* = 9.3, 3.0 Hz, 2H), 1.78 – 1.65 (m, 2H). $^{13}$C NMR (151 MHz, DMSO-$d_6$) *δ* 162.1, 142.7, 140.0, 136.1, 131.0, 129.1, 127.8, 124.2, 122.5, 120.9, 116.5, 113.6, 112.0, 44.8, 30.0, 12.5. $^{19}$F NMR (376 MHz, DMSO-$d_6$) *δ* 47.82, -74.61 (TFA). TLC $R_f$ = 0.23 (10% Methanol in DCM).

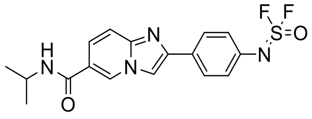

Exact Mass: 378.10
Molecular Weight: 378.40

(4-(6-(isopropylcarbamoyl)imidazo[1,2-*a*]pyridin-2-yl)phenyl)sulfurimidoyl difluoride (TFA salt, off-white solid, 5.5 mg). LRMS (+) calcd for $(M+H)^+$ 379.1. Found 379.1. $^1$H NMR (600 MHz, DMSO-$d_6$) δ 9.14 (s, 1H), 8.63 (s, 1H), 8.47 (d, $J$ = 7.6 Hz, 1H), 8.10 – 8.00 (m, 2H), 7.88 (d, $J$ = 9.5 Hz, 1H), 7.74 (d, $J$ = 9.4 Hz, 1H), 7.41 (d, $J$ = 8.5 Hz, 2H), 4.13 (h, $J$ = 6.7 Hz, 1H), 1.20 (d, $J$ = 6.6 Hz, 6H). $^{13}$C NMR (151 MHz, DMSO-$d_6$) δ 162.7, 158.2, 158.0, 135.4, 128.5, 127.5, 124.1, 116.8, 114.9, 114.5, 110.9, 41.3, 22.3. $^{19}$F NMR (376 MHz, DMSO-$d_6$) δ 47.85, -74.37 (TFA).

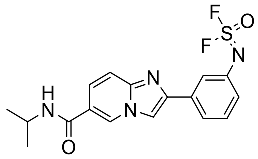

Exact Mass: 378.10
Molecular Weight: 378.40

(3-(6-(Isopropylcarbamoyl)imidazo[1,2-*a*]pyridin-2-yl)phenyl)sulfurimidoyl difluoride (TFA salt, off-white solid, 21.8 mg, 0.0449 mmol, 44% isolated yield). LRMS (+) calcd for $(M+H)^+$ 379.1, found 379.1. $^1$H NMR (600 MHz, DMSO-$d_6$) $\delta$ 9.17 (s, 1H), 8.70 (s, 1H), 8.51 (d, $J$ = 7.7 Hz, 1H), 7.94 (dd, $J$ = 9.4, 1.7 Hz, 1H), 7.91 (d, $J$ = 8.0 Hz, 1H), 7.85 (t, $J$ = 1.9 Hz, 1H), 7.79 (d, $J$ = 9.3 Hz, 1H), 7.59 (t, $J$ = 7.9 Hz, 1H), 7.31 (dd, $J$ = 8.0, 1.3 Hz, 1H), 4.13 (dq, $J$ = 13.5, 6.7 Hz, 1H), 1.20 (d, $J$ = 6.6 Hz, 6H). $^{13}$C NMR (151 MHz, DMSO-$d_6$) $\delta$ 162.5, 143.4, 140.8, 136.0, 132.8, 130.9, 128.7, 127.0, 124.1, 123.8, 122.1, 120.7, 114.3, 111.6, 41.4, 22.3. $^{19}$F NMR (376 MHz, DMSO-$d_6$) $\delta$ 49.55, -74.35 (TFA). TLC $R_f$ = 0.23 (10% Methanol in DCM).

**Representative procedure for the synthesis of sulfamide**

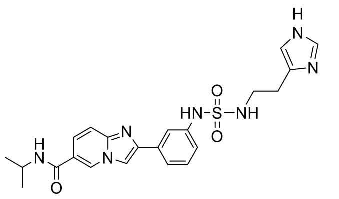

Exact Mass: 467.17
Molecular Weight: 467.55

To a DMSO solution of (3-(6-(isopropylcarbamoyl)imidazo[1,2-*a*]pyridin-2-yl)phenyl)sulfurimidoyl difluoride (~7 mg, 18.5 μmol) was added histamine (24 mg, 132 μmol) in PBS and stirred overnight at 37 °C. To this solution was added histamine (12 mg, 66 μmol) in PBS (pH adjusted to 10 by NaOH) and stirred overnight. The reaction mixture was directly purified on HPLC to give the target compound (7 mg, 14.6 μmol, 79%). LRMS (+) calcd for $(M+H)^+$ 468.2. Found 468.5. $^1$H NMR (600 MHz, DMSO-$d_6$) δ 9.86 (s, 1H), 9.11 (t, $J$ = 1.3 Hz, 1H), 8.91 (d, $J$ = 1.4 Hz, 1H), 8.46 (s, 1H), 8.40 (d, $J$ = 7.7 Hz, 1H), 7.81 – 7.73 (m, 2H), 7.69 (t, $J$ = 5.9 Hz, 1H), 7.65 (d, $J$ = 9.4 Hz, 1H), 7.60 (dt, $J$ = 7.7, 1.3 Hz, 1H), 7.37 (t, $J$ = 7.9 Hz, 1H), 7.31 (d, $J$ = 1.3 Hz, 1H), 7.18 – 7.12 (m, 1H), 4.17 – 4.07 (m, $J$ = 6.7 Hz, 1H), 3.19 (q, $J$ = 6.6 Hz, 2H), 2.80 (t, $J$ = 6.8 Hz, 2H), 1.20 (d, $J$ = 6.6 Hz, 6H). $^{13}$C NMR (151 MHz, DMSO-$d_6$) δ 163.0, 144.4, 144.3, 139.3, 133.7, 133.5, 130.5, 129.5, 128.4, 124.6, 120.8, 120.3, 118.2, 116.3, 115.7, 115.2, 110.4, 41.3, 40.8, 24.4, 22.3.

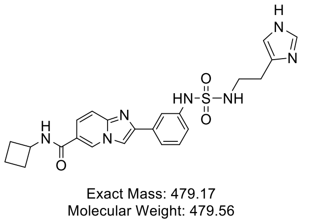


2-(3-((*N*-(2-(1H-imidazol-4-yl)ethyl)sulfamoyl)amino)phenyl)-*N*-cyclobutylimidazo[1,2-*a*]pyridine-6-carboxamide (TFA salt). LRMS (+) calcd for $(M+H)^+$ 480.2. Found 480.2. $^1$H NMR (600 MHz, DMSO-$d_6$) δ 9.86 (s, 1H), 9.12 (t, *J* = 1.4 Hz, 1H), 8.90 (d, *J* = 1.4 Hz, 1H), 8.79 (d, *J* = 7.4 Hz, 1H), 8.47 (s, 1H), 7.77 (p, *J* = 3.3 Hz, 2H), 7.74 – 7.65 (m, 2H), 7.60 (dt, *J* = 7.7, 1.3 Hz, 1H), 7.38 (d, *J* = 7.9 Hz, 1H), 7.31 (d, *J* = 1.3 Hz, 1H), 7.19 – 7.10 (m, 1H), 4.45 (h, *J* = 8.2 Hz, 1H), 3.19 (q, *J* = 6.7 Hz, 2H), 2.80 (t, *J* = 6.8 Hz, 2H), 2.25 (qt, *J* = 7.7, 2.8 Hz, 2H), 2.15 – 2.03 (m, 5H), 1.79 – 1.61 (m, 2H). $^{19}$F NMR (376 MHz, DMSO-$d_6$) δ -74.43. $^{13}$C NMR (151 MHz, DMSO-$d_6$) δ 162.7, 144.2, 143.9, 139.2, 133.6, 133.1, 130.3, 129.4, 128.4, 124.7, 120.5, 120.2, 118.2, 116.2, 115.6, 114.9, 110.4, 44.6, 40.7, 29.9, 24.3, 14.7.

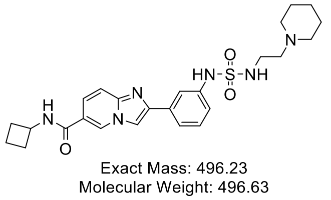


*N*-cyclobutyl-2-(3-((N-(2-(piperidin-1-yl)ethyl)sulfamoyl)amino)phenyl)imidazo[1,2-*a*]pyridine-6-carboxamide. LRMS (+) calcd for $(M+H)^+$ 497.2. Found 497.3.$^1$H NMR (600 MHz, DMSO-$d_6$) δ 9.07 (dd, *J* = 1.8, 1.0 Hz, 1H), 8.73 (d, *J* = 7.4 Hz, 1H), 8.44 (d, *J* = 0.7 Hz, 1H), 8.28 – 8.19 (m, 2H), 7.82 (t, *J* = 1.9 Hz, 1H), 7.68 (dd, *J* = 9.5, 1.8 Hz, 1H), 7.63 – 7.57 (m, 2H), 7.36 (t, *J* = 7.9 Hz, 1H), 7.22 (s, 1H), 7.16 (ddd, *J* = 8.0, 2.3, 1.0 Hz, 1H), 4.46 – 4.42 (m, 1H), 2.96 (t, *J* = 7.0 Hz, 2H), 2.28 (dd, *J* = 11.3, 4.5 Hz, 2H), 2.27 – 2.16 (m, 6H), 2.13 – 2.10 (m, 2H), 1.75 – 1.65 (m, 2H), 1.37 – 1.34 (m, 4H), 1.30 – 1.21 (m, 2H). $^{13}$C NMR (151 MHz, DMSO-$d_6$) δ 163.1, 145.5, 144.9, 139.4, 134.3, 129.3, 128.2, 123.6, 120.1, 119.9, 118.0, 115.7, 115.6, 110.2, 68.5, 57.2, 55.8, 53.9, 44.6, 25.4, 23.9, 14.8.

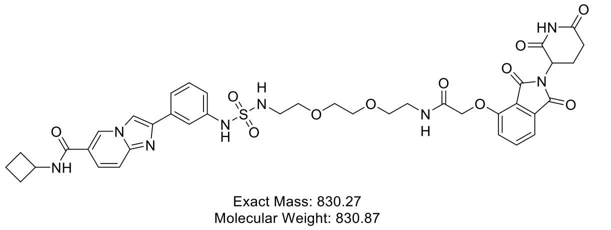


*N*-cyclobutyl-2-(3-((N-(2-(2-(2-(2-((2-(2,6-dioxopiperidin-3-yl)-1,3-dioxoisoindolin-4-yl)oxy)acetamido)ethoxy)ethoxy)ethyl)sulfamoyl)amino)phenyl)imidazo[1,2-*a*]pyridine-6-carboxamide. LRMS (+) calcd for $(M+H)^+$ 831.3. Found 831.4. $^1$H NMR (600 MHz, Methanol-$d_4$) δ 9.11 (dd, *J* = 1.7, 1.0 Hz, 1H), 8.45 (d, *J* = 0.7 Hz, 1H), 8.07 (dd, *J* = 9.4, 1.7 Hz, 1H), 7.78 (dt, *J* = 9.4, 0.9 Hz, 1H), 7.78 – 7.69 (m, 2H), 7.53 (ddd, *J* = 7.7, 1.8, 1.0 Hz, 1H), 7.46 – 7.35 (m, 2H), 7.32 (dd, *J* = 8.5, 0.6 Hz, 1H), 7.25 (ddd, *J* = 8.1, 2.2, 1.0 Hz, 1H), 5.09 (dd, *J* = 12.8, 5.5 Hz, 1H), 4.65 (d, *J* = 1.6 Hz, 2H), 4.55 – 4.47 (m, 1H), 3.56 (s, 1H), 3.58 – 3.49 (m, 8H), 3.39 (dd, *J* = 5.9, 4.8 Hz, 2H), 3.36 – 3.30 (m, 1H), 3.21 – 3.14 (m, 2H), 2.86 (ddd, *J* = 17.4, 13.9, 5.4 Hz, 1H), 2.77 – 2.61 (m, 2H), 2.42 – 2.34 (m, 2H), 2.21 – 2.07 (m, 3H), 1.84 (s, 1H), 1.87 – 1.76 (m, 2H). $^{13}$C NMR (151 MHz, Methanol-$d_4$) δ 174.6, 171.5, 169.9, 168.2, 167.5, 164.7, 156.0, 144.0, 141.7, 141.2, 138.2, 134.7, 131.3, 130.7, 130.4, 130.4, 125.1, 122.0, 121.8, 121.4, 119.0, 117.8, 117.5, 114.1, 112.8, 71.3, 71.2, 70.8, 70.2, 69.0, 56.1, 43.9, 40.1, 31.24, 31.22, 29.5, 23.6, 16.1.

**Safety statement.**

No unexpected or unusually high safety hazards were encountered.

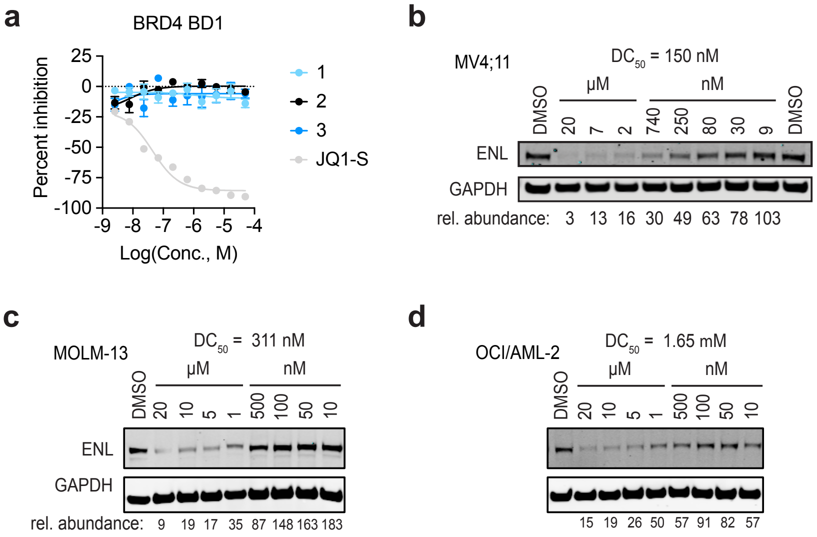


**Figure S1. Characterization of amido-imidazopyridine scaffold.** (a) Activity against BRD4 BD1 measured by HTRF. Mean ± s.e.m., $n = 4$. (b) immunoblot of ENL and GAPDH signal after 4h treatment with SR-1114 in Mv4;11. Relative abundance given as the ratio of ENL over GAPDH signal adjusted to percent of DMSO control. Half maximal effective degradation concentration ($DC_{50}$) determined by least squares fit. (c, d) as in (b) but with the indicated cell lines.

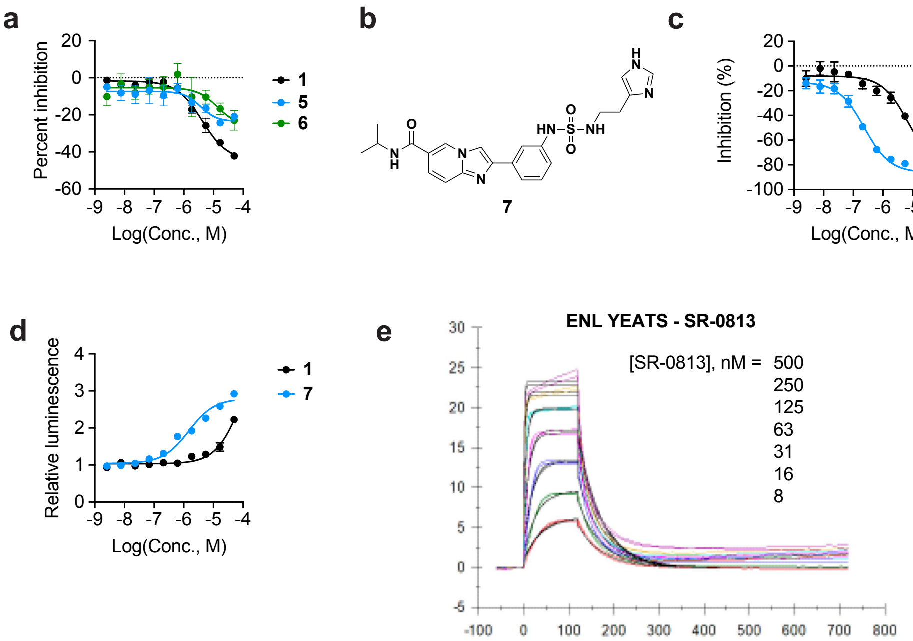


**Figure S2. Evaluation of SuFEx-based medicinal chemistry precursors and hits. (**a) Activity against the ENL YEATS domain measured by HTRF. Mean ± s.e.m., *n* = 2. (b) Chemical structure of Compound 7, a hit from SuFEx-based medicinal chemistry screen. (c) Activity against the ENL YEATS domain measured by HTRF. Mean ± s.e.m., *n* = 4. (d) Engagement of ENL(YEATS)-HiBiT measured by ligand-induced luminescence. Signal normalized to DMSO ± s.e.m., *n* = 4. (e) SPR sensorgram of ENL YEATS and SR-0813 binding data presented in Fig. 2g.

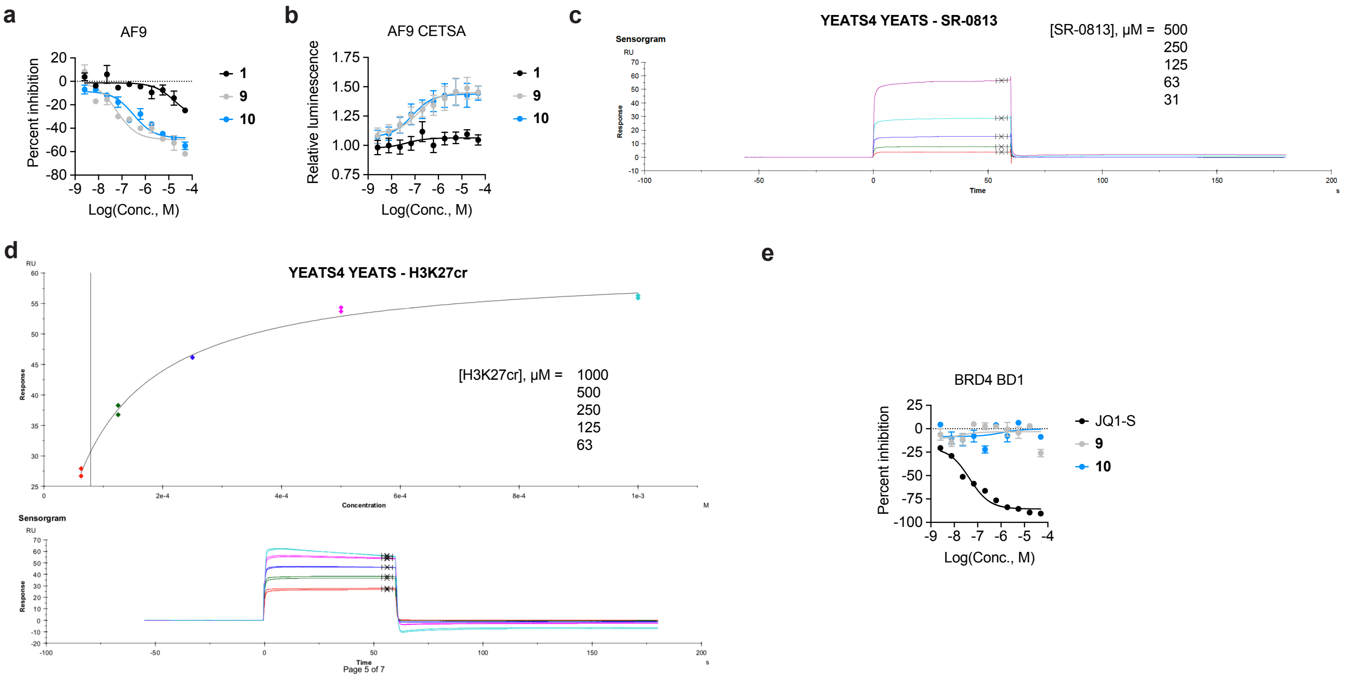


**Figure S3. Profiling SR-0813 against other acyl-lysine reader domains.** (a) Activity against the AF9 YEATS domain measured by HTRF. Mean ± s.e.m., *n* = 4. (b) Engagement of AF9(YEATS)-HiBiT measured by ligand-induced luminescence. Signal normalized to DMSO ± s.e.m., *n* = 4. (c) SPR sensorgram of YEATS4 YEATS and SR-0813 binding data presented in Fig. 2g. (d) Binding data and SPR sensorgram of YEATS4 YEATS and a peptide substrate H3K27cr. Used as a positive control to ensure proper folding of YEATS4 YEATS with expected binding to H3K27cr. (e) Activity against the BRD4 BD1 measured by HTRF. Mean ± s.e.m., *n* = 4.

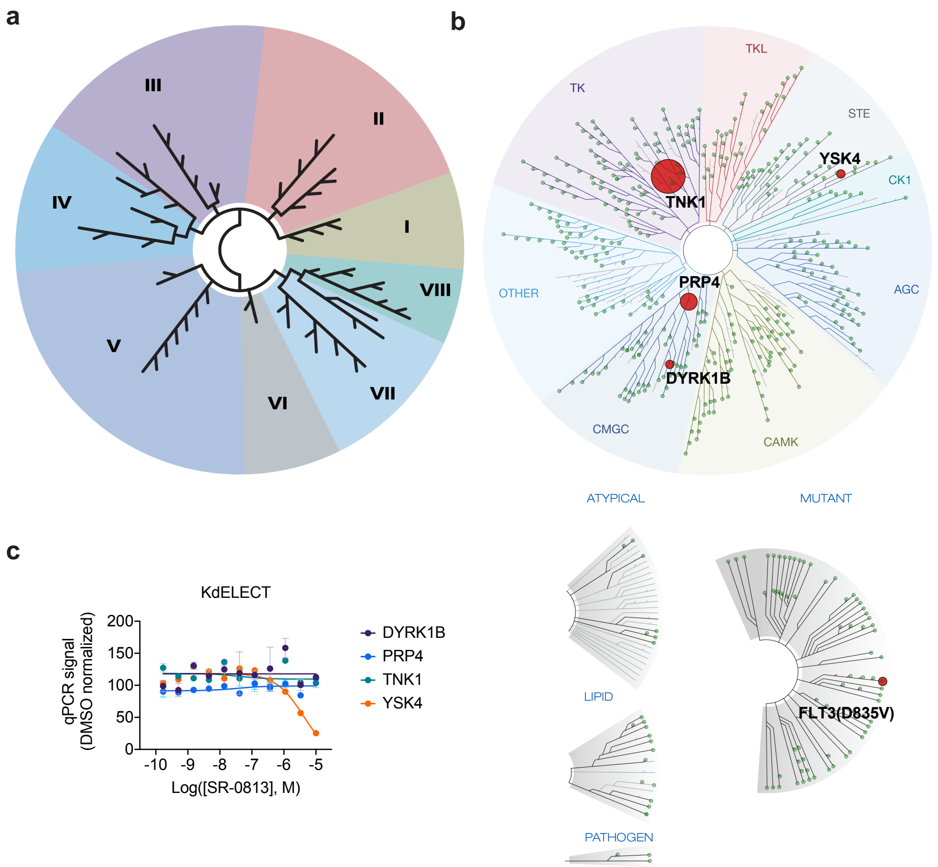


**Figure S4. Profiling SR-0813 for bromodomain and kinase off targets.** (a) BROMO*scan* profiling service revealed no binding by SR-0813 to any of 32 bromodomains tested. (b) KINOME*scan* profiling service (scanMAX) testing activity of SR-0813 against 468 kinase targets. (c) Dose-response evaluation of scanMAX hits in (b).

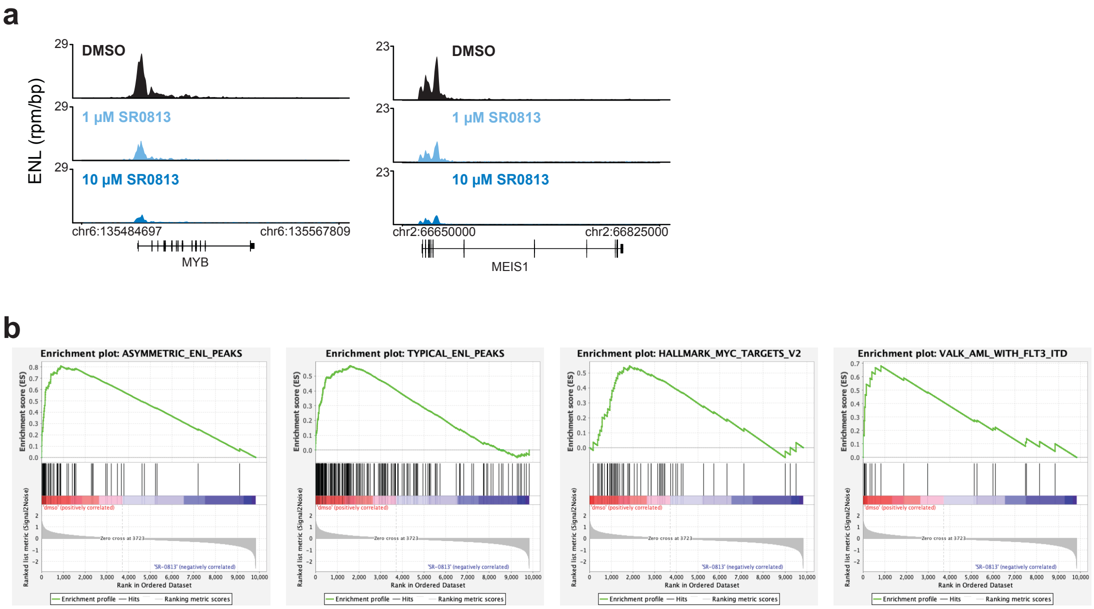


**Figure S5. SR-08-13 effects on ENL-dependent transcription.** (a) Gene tracks of ChIP-seq signal at asymmetrically loaded loci. (b) GSEA plots of top 4 enriched gene sets in Fig. 4f.
