## Supplemental Table 2 for "Chemical inhibition of ENL/AF9 YEATS domains in acute leukemia"

| plate# | well | CAS | Structure | MW | Inhibition %  Amine only | Inhibition %  Meta  2µM final  Rep1 | Inhibition %  Meta  2µM final  Rep2 | Inhibition %  Para  2µM final  Rep1 | Inhibition %  Para  2µM final  Rep2 |
| --- | --- | --- | --- | --- | --- | --- | --- | --- | --- |
| 1 | A1 | 2978-58-7 | 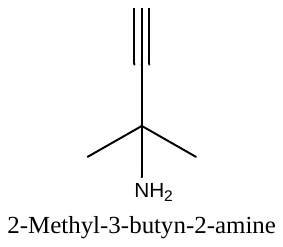 | 83 | -11.3 | -52.2 | -42.4 | -39.4 | -35.6 |
| 1 | A2 | 929-06-6 | 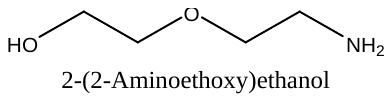 | 105.14 | 0.1 | -89.3 | -73.2 | -36.4 | -30.2 |
| 1 | A3 | 2906.12.09 | 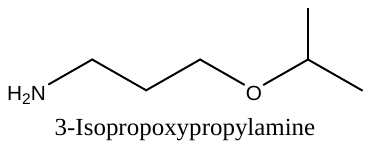 | 117.19 | -1.6 | -62.5 | -46.5 | -27.8 | -24.4 |
| 1 | A4 | 13325-10-5 | 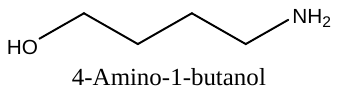 | 89 | 37.0 | -85.5 | -76.7 | -22.7 | -22.0 |
| 1 | A5 | 1003-03-8 | 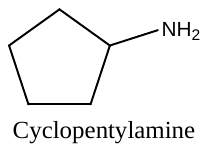 | 85.15 | 1.7 | -56.2 | -49.3 | -21.6 | -17.6 |
| 1 | A6 | 109-76-2 | 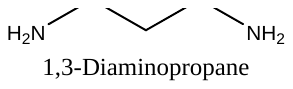 | 74.12 | 16.8 | -76.8 | -67.3 | -34.2 | -33.1 |
| 1 | A7 | 109-73-9 | 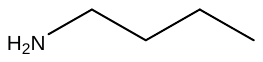 | 73 | 4.5 | -46.7 | -38.5 | -21.9 | -25.3 |
| 1 | A8 | 156-87-6 | 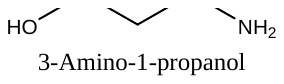 | 75 | -8.3 | -90.8 | -71.2 | -35.2 | -39.0 |
| 1 | A9 | 6291-84-5 | 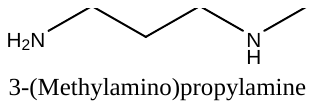 | 88 | -1.7 | -73.4 | -67.9 | -16.5 | -11.8 |
| 1 | A10 | 2867-59-6 | 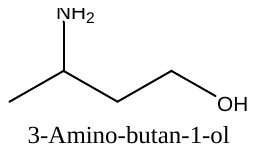 | 89 | 4.2 | -57.9 | -47.0 | -15.3 | -16.2 |
| 1 | B1 | 115-70-8 | 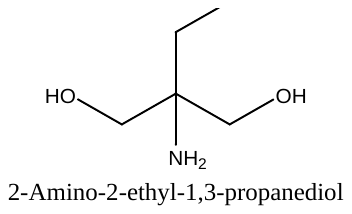 | 119 | -9.1 | -66.9 | -56.5 | -28.0 | -27.0 |
| 1 | B2 | 87120-72-7 | 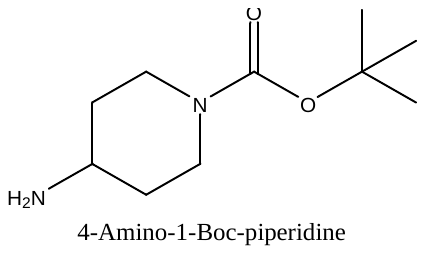 | 200 | 24.5 | -71.4 | -64.4 | -27.9 | -24.5 |
| 1 | B3 | 60142-96-3 | 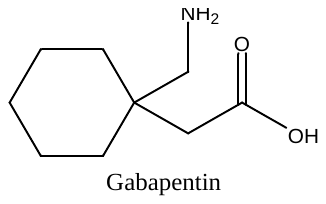 | 171 | -0.4 | -49.2 | -39.7 | -27.9 | -24.7 |
| 1 | B4 | 120-20-7 | 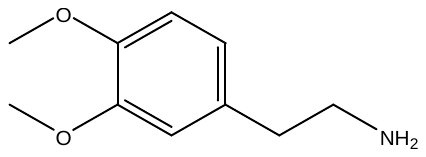 | 181 | 0.3 | -58.7 | -46.9 | -15.5 | -16.9 |
| 1 | B5 | 3731-52-0 | 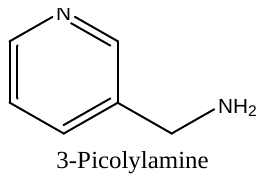 | 108 | -2.9 | -77.0 | -67.7 | -31.1 | -32.7 |
| 1 | B6 | 3300-51-4 | 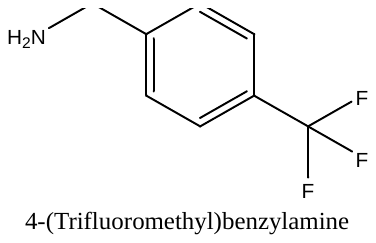 | 175 | 1.3 | -38.5 | -29.4 | -16.0 | -22.4 |
| 1 | B7 | 26177-43-5 | 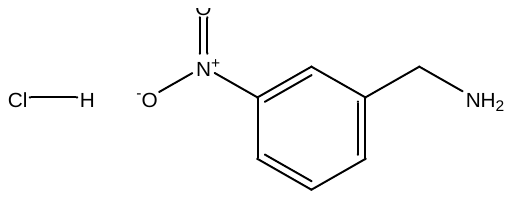 | 189 | 3.5 | -69.5 | -58.2 | -19.1 | -17.3 |
| 1 | B8 | 1118-89-4 | 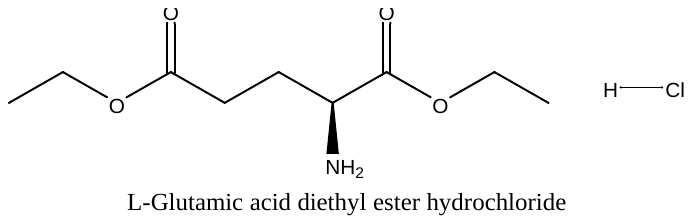 | 240 | -1.1 | -49.7 | -45.6 | -23.9 | -15.5 |
| 1 | B9 | 123-00-2 | 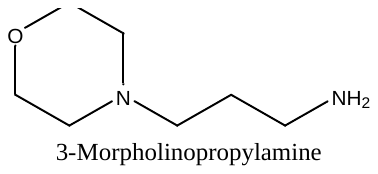 | 144 | 2.3 | -75.9 | -71.4 | -23.1 | -22.7 |
| 1 | B10 | 156917-23-6 | 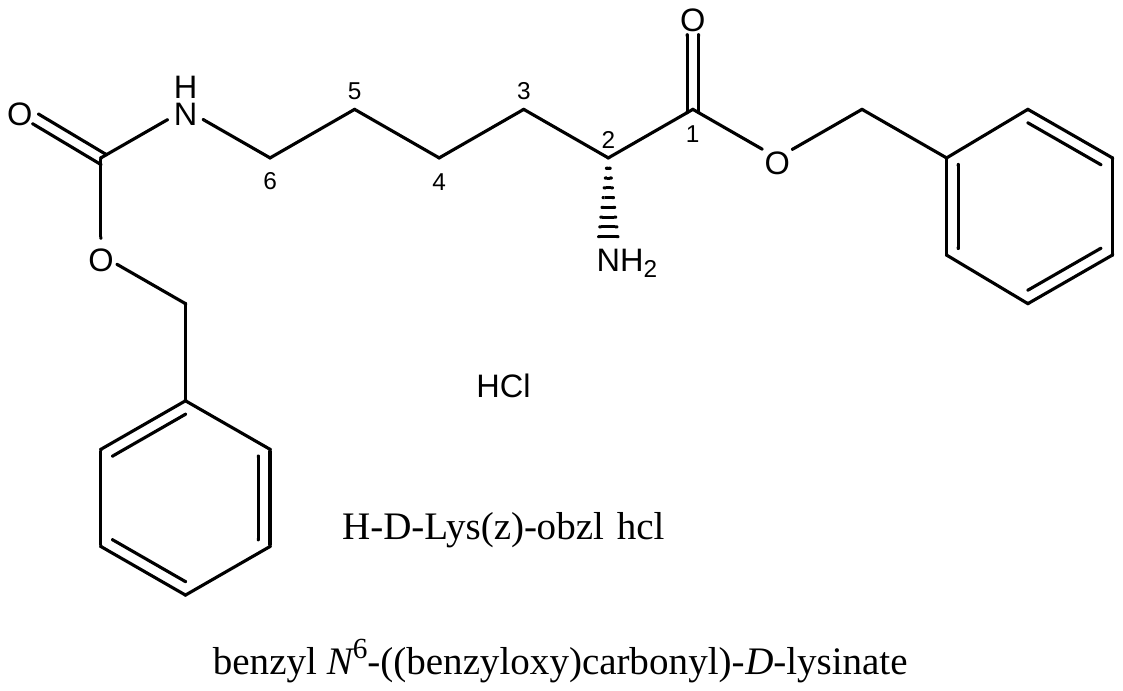 | 407 | 3.0 | -54.6 | -57.7 | -13.2 | -19.5 |
| 1 | C1 | 57260-73-8 | 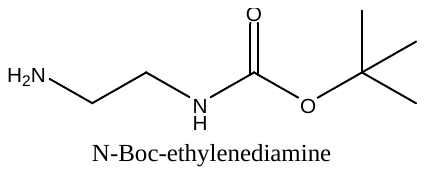 | 160 | 6.4 | -75.0 | -66.0 | -18.4 | -16.4 |
| 1 | C2 | 439117-39-2 | 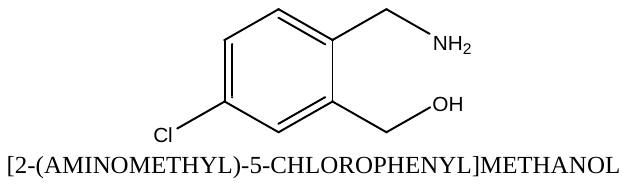 | 172 | 0.4 | -64.0 | -59.9 | -19.7 | -34.4 |
| 1 | C3 | 2491-18-1 | 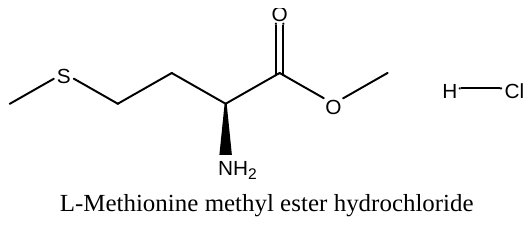 | 200 | 0.9 | -57.0 | -44.7 | -24.8 | -17.1 |
| 1 | C4 | 56-92-8 | 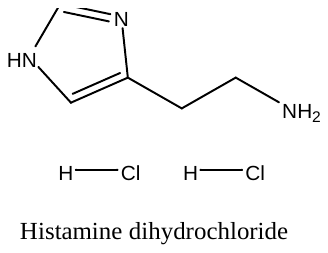 | 184 | 3.2 | -93.4 | -83.0 | -31.6 | -17.7 |
| 1 | C5 | 696-60-6 | 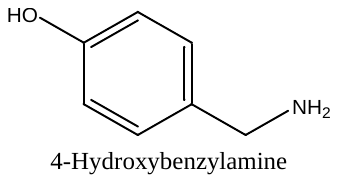 | 123 | 4.1 | -89.2 | -74.0 | -26.6 | -23.0 |
| 1 | C6 | 100-46-9 | 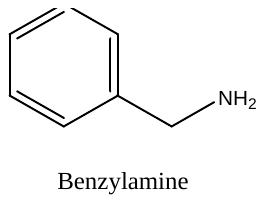 | 107 | 4.9 | -79.8 | -68.2 | -17.1 | -11.2 |
| 1 | C7 | 63649-14-9 | 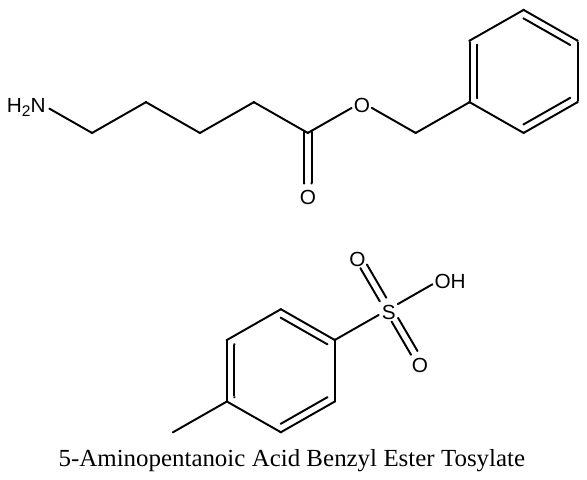 | 379 | 8.1 | -37.1 | -28.0 | 3.0 | -3.2 |
| 1 | C8 | 22572-33-4 | 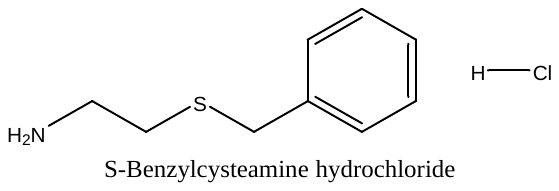 | 204 | -0.6 | -52.2 | -44.4 | -23.8 | -18.5 |
| 1 | C9 | 2039-66-9 | 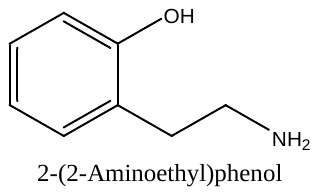 | 137.18 | 2.1 | -21.8 | -69.8 | -30.5 | -23.5 |
| 1 | C10 | 35303-76-5 | 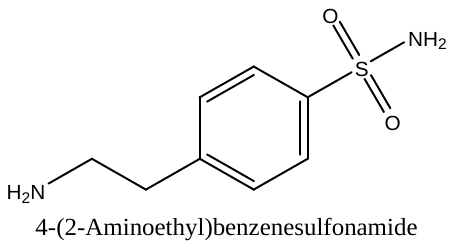 | 200 | 12.2 | -86.1 | -72.2 | -19.5 | -18.5 |
| 1 | D1 | 16652-64-5 | 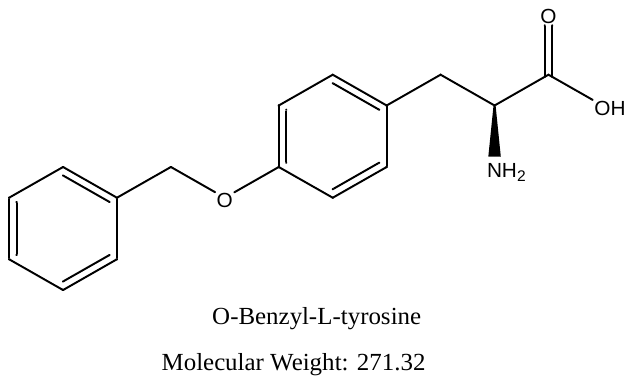 | 271 | 18.5 | -41.6 | -26.7 | -11.5 | -15.0 |
| 1 | D2 | 3417-91-2 | 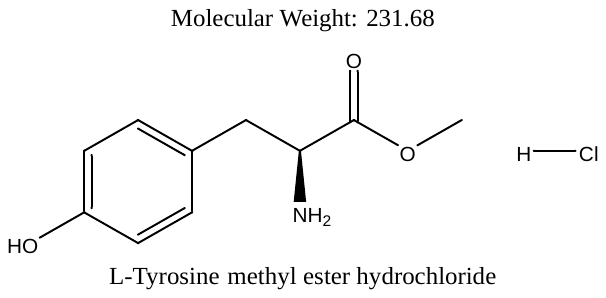  Phenol can react as well | 232 | -0.5 | -50.5 | -36.7 | -19.7 | -10.3 |
| 1 | D3 | 60-32-2 | 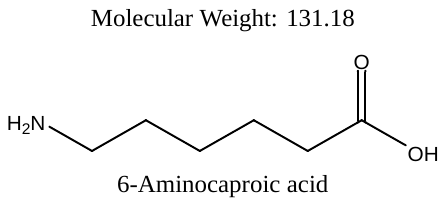 | 131 | -2.7 | -59.7 | -51.9 | -16.7 | -16.3 |
| 1 | D4 | 57213-48-6 | 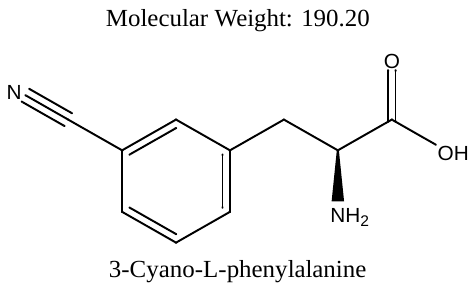 | 190 | -0.8 | -45.6 | -33.0 | -19.3 | -15.9 |
| 1 | D5 | 5466-22-8 | 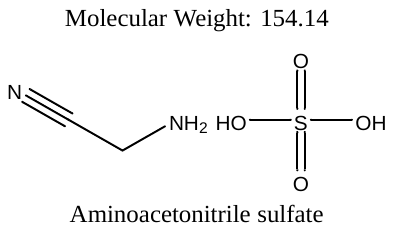 | 154 | 7.6 | -55.4 | -47.1 | -15.4 | -8.6 |
| 1 | D6 | 19883-74-0 | 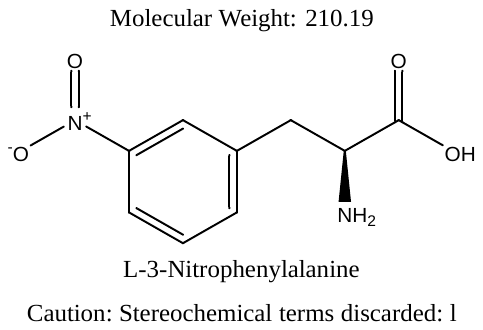 | 210 | 1.0 | -48.8 | -34.9 | -16.3 | -13.7 |
| 1 | D7 | 4083-572 | 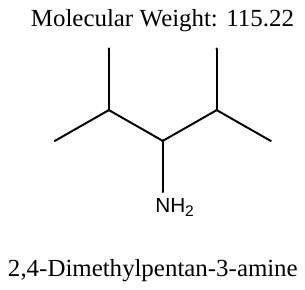 | 115 | -2.4 | -60.5 | -43.8 | -20.5 | -14.8 |
| 1 | D8 |  | 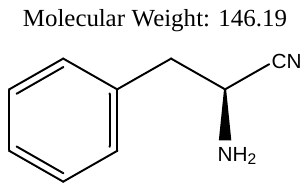 | 146 | 1.4 | -48.7 | -38.1 | -23.1 | -16.4 |
| 1 | D9 | 14464-68-7 | 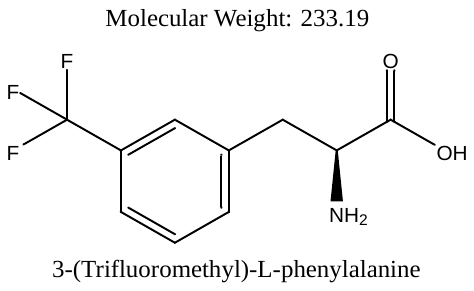 | 233 | 0.4 | -42.2 | -27.7 | 2.6 | 24.5 |
| 1 | D10 | 80126-51-8 | 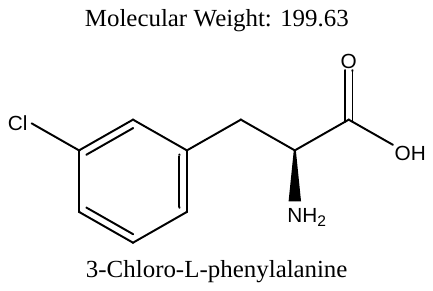 | 199 | -1.9 | -42.1 | -29.2 | -13.7 | -6.1 |
| 1 | E1 | 19883-77-3 | 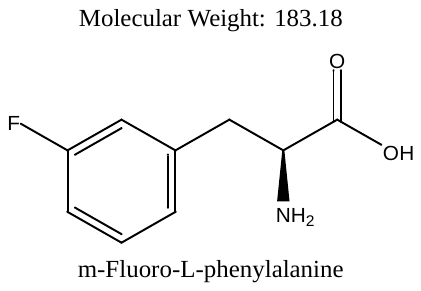 | 183 | -0.7 | -30.6 | -16.2 | -13.9 | -11.2 |
| 1 | E2 | 103616-89-3 | 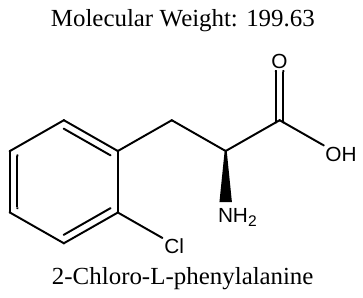 | 199 | 31.5 | -23.4 | -14.8 | -5.4 | -3.2 |
| 1 | E3 | 72-18-4 | 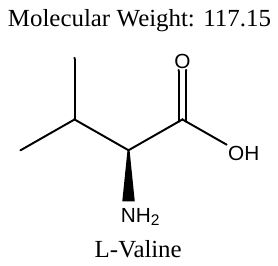 | 117 | 6.6 | -39.5 | -18.7 | -2.1 | 2.1 |
| 1 | E4 | 3182-93-2 | 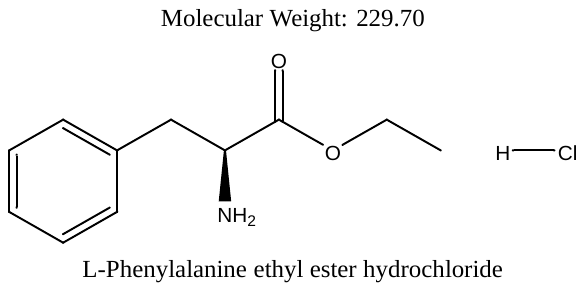 | 230 | 3.4 | -54.3 | -59.6 | -24.5 | -19.0 |
| 1 | E5 | 28211-04-3 | Poly epsilon L-lysine HCl | 385 | 49.7 | -54.7 | -41.9 | -25.0 | -20.8 |
| 1 | E6 | 6850-28-8 | 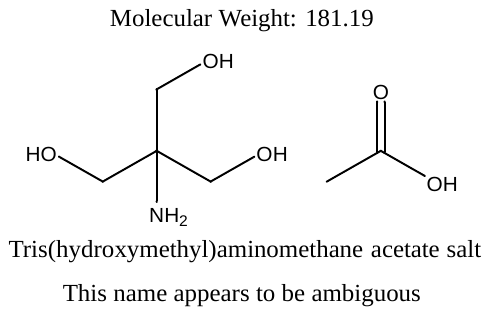 | 181 | 20.7 | -40.1 | -26.1 | 1.5 | 10.4 |
| 1 | E7 | 3048/01/09 | 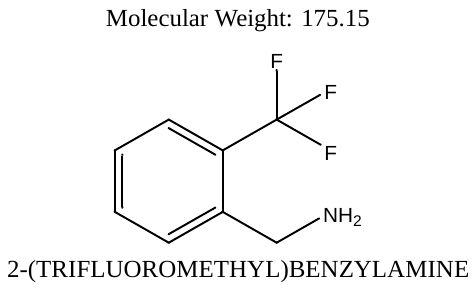 | 175 | 119.9 | -46.4 | -35.5 | -15.5 | -8.3 |
| 1 | E8 | 107-10-8 | 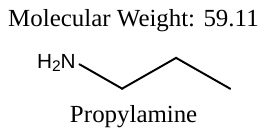 | 59 | 2.9 | -48.4 | -28.9 | -10.5 | -11.3 |
|  | E9 |  |  |  | 30.2 | -42.7 | -28.6 | -9.1 | -7.0 |
|  | E10 |  |  |  | 3.4 | -44.2 | -32.4 | -20.6 | -15.5 |
| 1 | F1 | 116668-47-4 | 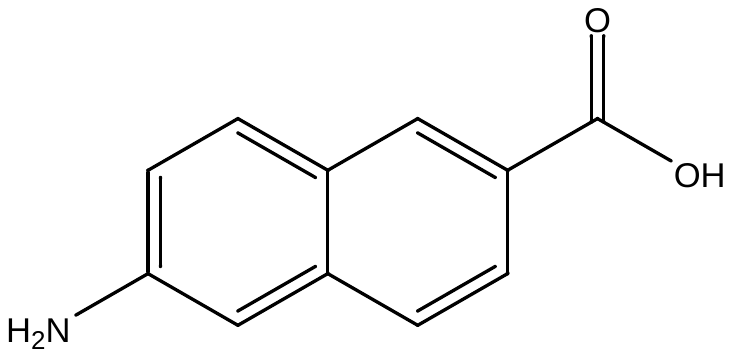 | 187 | -6.4 | -54.5 | -38.5 | -26.8 | -21.9 |
| 1 | F2 | 4214-76-0 | 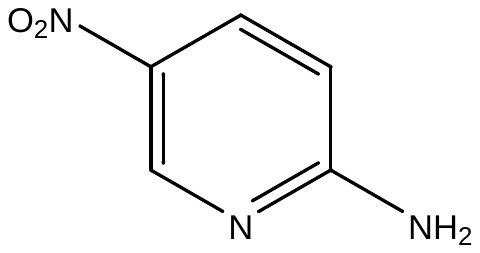 | 139 | -6.4 | -45.7 | -38.9 | -14.0 | -10.3 |
| 1 | F3 | 5978-75-6 | 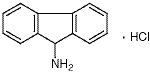 | 217.7 | 4.4 | -48.8 | -41.6 | -20.7 | -14.9 |
| 1 | F4 | 123-30-8 | 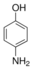 | 109 | 99.9 | 23.1 | 20.1 | 50.3 | 45.1 |
| 1 | F5 | 61437-85-2 | 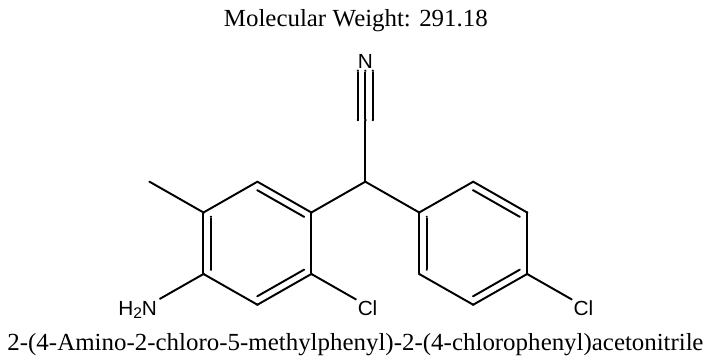 | 291 | 0.2 | -53.7 | -43.2 | -20.0 | -15.0 |
| 1 | F6 | 99-31-0 | 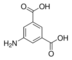 | 181 | 4.2 | -47.9 | -33.6 | -8.3 | -9.1 |
| 1 | F7 | 30433-91-1 | 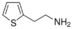 | 127 | 24.6 | -87.1 | -78.3 | -32.3 | -26.1 |
| 1 | F8 | 4122/04/07 | 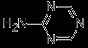 | 96 | -1.1 | -48.5 | -40.0 | -18.0 | -14.0 |
| 1 | F9 | 75-64-9 | 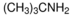 | 73 | -5.4 | -46.0 | -31.9 | -21.5 | -14.7 |
| 1 | F10 | 108607-02-9 | 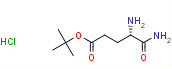 | 239 | -4.9 | -46.4 | -40.7 | -25.2 | -20.3 |
| 1 | G1 | 100-82-3 | 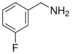 | 125 | -13.3 | -92.9 | -86.6 | -34.9 | -42.4 |
| 1 | G2 | 20781-21-9 | 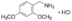 | 204 | -6.0 | -42.4 | -36.2 | -15.7 | -6.5 |
| 1 | G3 | 132388-58-0 | 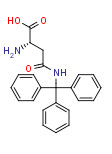 | 374 | -0.3 | -54.8 | -36.4 | -18.1 | -12.9 |
| 1 | G4 | 2393-23-9 | 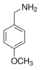 | 137 | 4.3 | -73.5 | -63.3 | -17.0 | -21.0 |
| 1 | G5 | 140-75-0 | 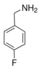 | 125 | 7.1 | -82.0 | -73.4 | -33.4 | -29.1 |
| 1 | G6 | 122-80-5 | 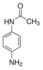 | 150 | -6.1 | -45.5 | -36.8 | -26.5 | -26.0 |
| 1 | G7 | 593-51-1 | 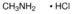 | 68 | 3.2 | -83.9 | -76.6 | -19.1 | -21.1 |
| 1 | G8 | 20859-02-3 L-tert-Leucine | 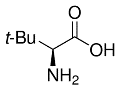 | 131 | -3.0 | -59.3 | -44.4 | -50.4 | -31.8 |
| 1 | G9 | 04-12-5198 | 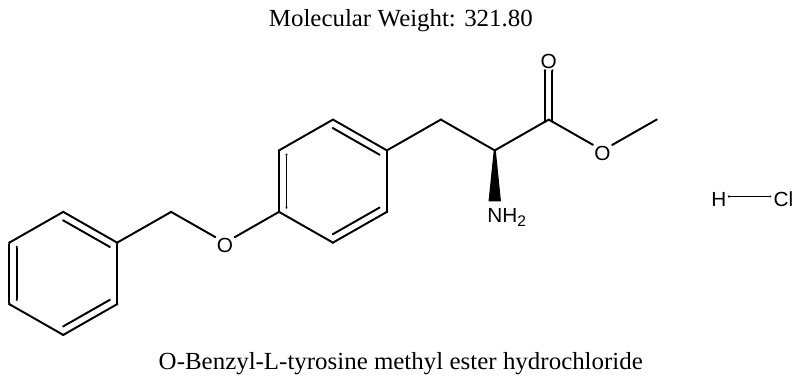 | 322 | 53.4 | -30.8 | -5.7 | -34.9 | 4.4 |
| 1 | G10 | 1798-50-1 | 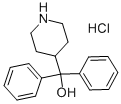 | 304 | 3.7 | -22.2 | -20.6 | -7.7 | -5.1 |
| 2 | A1 | 32462-30-9 | 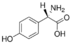 | 167 | -11.1 | -35.0 | -24.4 | 7.6 | -11.1 |
| 2 | A2 | 3490/06/06 | 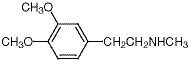 | 195 | 9.4 | -38.6 | -31.0 | -8.2 | -8.8 |
| 2 | A3 | 18542-42-2 | 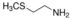 | 91 | -11.4 | -73.2 | -67.1 | -16.9 | -29.3 |
| 2 | A4 | 459-19-8 | 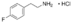 | 176 | 4.6 | -53.9 | -39.8 | -20.2 | -10.1 |
| 2 | A5 | Bis(3-aminopropyl)ether | 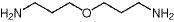 | 132 | 0.0 | -77.3 | -68.2 | -20.5 | -15.5 |
| 2 | A6 | 150517-77-4 | 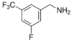 | 193 | -6.9 | 71.6 | -36.7 | -29.7 | -16.0 |
| 2 | A7 | 15996-76-6 | 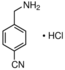 | 169 | 1.3 | -70.8 | -58.4 | -36.1 | -29.6 |
| 2 | A8 | 104-86-9 | 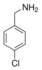 | 162 | -18.4 | -36.3 | -27.5 | -32.0 | -23.3 |
| 2 | A9 | 7663-77-6 | 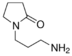 | 142 | -14.6 | -75.8 | -71.6 | -57.2 | -30.8 |
| 2 | A10 | 156-41-2 | 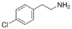 | 156 | 58.6 | -38.7 | -35.5 | -36.1 | -34.7 |
| 2 | B1 | 3886-70-2 | 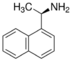 | 171 | -2.8 | -45.8 | -42.5 | -20.3 | -11.7 |
| 2 | B2 | 696-40-2 | 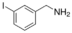 | 233 | 7.3 | -29.8 | -19.6 | -0.6 | -0.1 |
| 2 | B3 | 2432-99-7 | 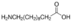 | 201 | -2.3 | -36.2 | -22.2 | -13.0 | -13.3 |
| 2 | B4 | 70-78-0 | 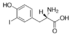 | 307 | -7.6 | -53.3 | -23.5 | -17.3 | -14.9 |
| 2 | B5 | 492-41-1 | 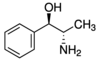 | 151 | -11.2 | -57.9 | -51.3 | -34.4 | -30.0 |
| 2 | B6 | 4747-21-1 | 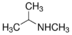 | 73 | -0.5 | -47.6 | -38.3 | -19.8 | -18.9 |
| 2 | B7 | 2017-67-6 | 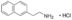 | 208 | 30.7 | -26.0 | 9.7 | -8.1 | -9.0 |
| 2 | B8 | 78-81-9 | 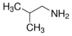 | 73 | -3.4 | -55.3 | -41.9 | -28.1 | -18.6 |
| 2 | B9 | 2935-35-5 | 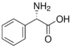 | 151 | -11.6 | -62.8 | -78.0 | -27.5 | -21.7 |
| 2 | B10 | DL-4-hydroxyphenylglycine | 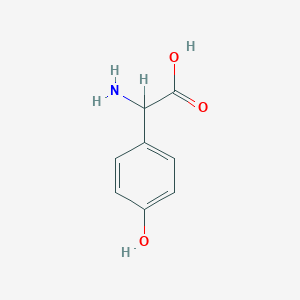 | 167 | -0.3 | -53.3 | -46.5 | -17.0 | -24.1 |
| 2 | C1 | 4104-45-4 | 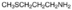 | 105 | 29.4 | -82.9 | -67.7 | -16.6 | -19.8 |
| 2 | C2 | 1986-47-6 | 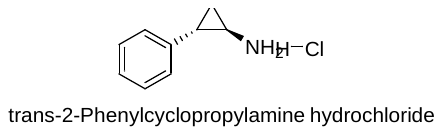 | 170 | 1.9 | -36.1 | -26.1 | -21.4 | -16.4 |
| 2 | C3 | 04-12-5147 | 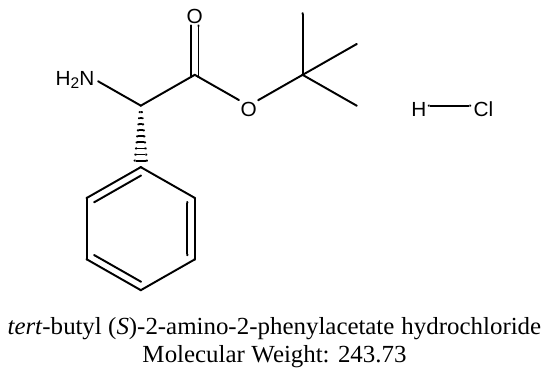 | 244 | -48.2 | -63.8 | -51.3 | -33.0 | -27.2 |
| 2 | C4 | 150-30-1 | 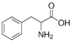 | 165 | 4.4 | -44.6 | -31.4 | -23.6 | -15.1 |
| 2 | C5 | H-Asp(otBu)-OtBu/HCl | 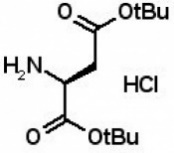 | 282 | -5.7 | -51.5 | -40.9 | -14.5 | -10.3 |
| 2 | C6 | 13288-57-8 | 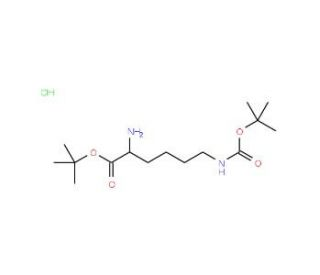 | 339 | 0.7 | -48.8 | -21.3 | -17.1 | -20.0 |
| 2 | C7 | H-His(Trt)-OtBu/HCl | 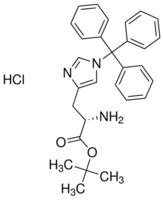 | 490 | 191.2 | -50.2 | -41.9 | -14.3 | -13.8 |
| 2 | C8 | 16874-09-2 | 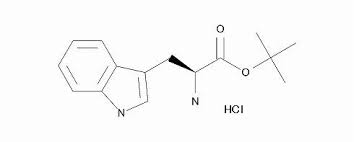 | 297 | 3.1 | -56.3 | -43.1 | -14.6 | -7.6 |
| 2 | C9 | H-Ser-OtBu/HCl | 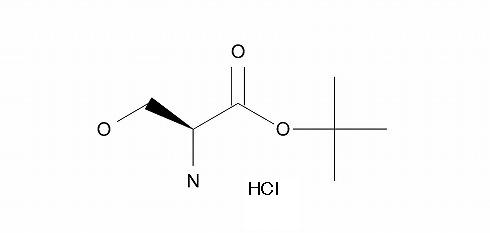 | 198 | 5.3 | -64.8 | -52.4 | -19.6 | -17.3 |
| 2 | C10 | 04-10-0073 | D-cystein 349-46-2 | 240 | 0.0 | -50.7 | -44.4 | -16.9 | -15.3 |
| 2 | D1 | 04-12-5047 | 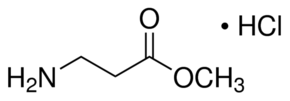 | 103 | 2.6 | -77.3 | -76.9 | -23.1 | -21.3 |
| 2 | D2 | 04-10-0016 | D-glutamic acid 6893-26-1 | 147 | 8.1 | -26.3 | -14.9 | -10.7 | -1.8 |
| 2 | D3 | H-Asn-OH | 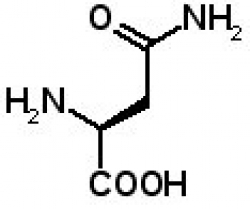 | 132 | 28.7 | -47.8 | -35.7 | -16.4 | -9.4 |
| 2 | D4 | 73-32-5 | 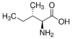 | 131 | -4.7 | -44.7 | -42.4 | -23.7 | -23.0 |
| 2 | D5 | 27894-50-4 | 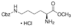 | 331 | -9.8 | -67.8 | -56.9 | -23.1 | -21.3 |
| 2 | D6 | 51537-21-4 | 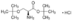 | 253 | 52.2 | -49.7 | -49.6 | -21.3 | -15.2 |
| 2 | D7 | 18905-73-2 | 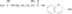 | 336 | -6.4 | -51.7 | -39.3 | -24.1 | -20.1 |
| 2 | D8 | 14907-27-8 | 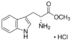 | 255 | -2.8 | -80.1 | -72.8 | -27.8 | -23.3 |
| 2 | D9 | 04-13-5043 | 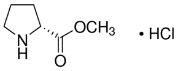 | 166 | -1.3 | -76.4 | -70.3 | -42.1 | -33.6 |
| 2 | D10 | L-Valyl-L-Leucine | 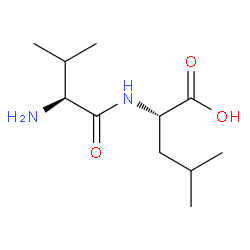 | 230 | -3.3 | -57.1 | -37.8 | -18.9 | -15.5 |
| 2 | E1 | H-a-Abu-otBu/HCl | 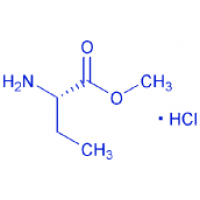 | 154 | 27.4 | -71.2 | -55.9 | -15.0 | -10.1 |
| 2 | E2 | H-D-Cys(tBu)-OH/HCl | 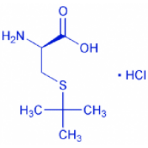 | 214 | 1.4 | -41.5 | -33.0 | -13.3 | -2.9 |
| 2 | E3 | 04-12-5044 | 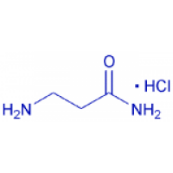 | 125 | -1.5 | -100.0 | -104.0 | -29.7 | -25.3 |
| 2 | E4 | H-D-Phe(3-Ome)-OH | 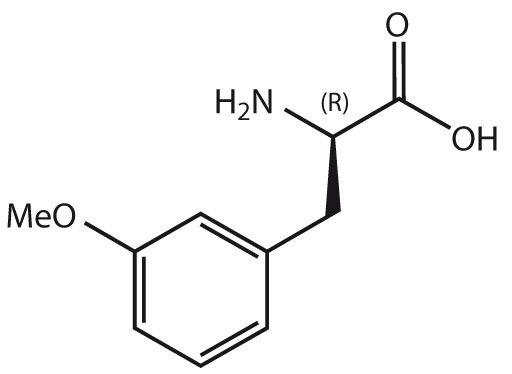 | 195 | 2.2 | -39.8 | -29.0 | -21.0 | -16.2 |
| 2 | E5 | H-D-Asp(Ome)-OH/ | 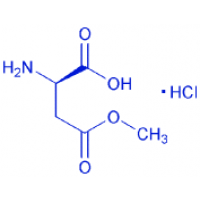 | 184 | -26.1 | -63.5 | -52.7 | -20.6 | -34.1 |
| 2 | E6 | 13188-89-1 | 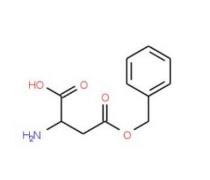 | 223 | 21.9 | -51.3 | -27.8 | -24.0 | -18.8 |
| 2 | E7 | 75/04/07 | 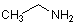  ca. 10% in Tetrahydrofuran, ca. 2mol/L?? | 45 | -8.1 | -40.7 | -25.8 | -11.3 | -9.2 |
| 2 | E8 | 141-43-5 | 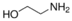 | 61 | -2.0 | -95.3 | -88.1 | -32.4 | -35.6 |
| 2 | E9 | 04-12-5196 | H-Tyr(Bzl)-Obzl/HCl | 398 | 3.0 | -54.5 | -62.6 | -16.0 | -11.2 |
| 2 | E10 | 04-12-5217 | 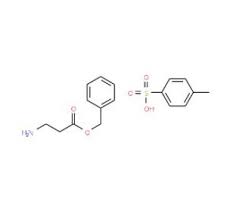 | 351 | 16.6 | -63.7 | -60.3 | -3.2 | -0.7 |
| 2 | F1 | 69320-89-4 | 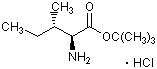 | 224 | -2.4 | -40.4 | -34.8 | -17.1 | -9.6 |
| 2 | F2 | 04-12-5145 | 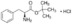 | 258 | 4.7 | -48.8 | -38.4 | -23.6 | -14.0 |
| 2 | F3 | 04-12-5075 | 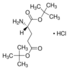 | 296 | -1.3 | -57.4 | -43.2 | -16.6 | -10.6 |
| 2 | F4 | 04-12-5087 | 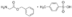 | 337 | 5.0 | -63.5 | -60.4 | -23.4 | -16.4 |
| 2 | F5 | 32677-01-3 | 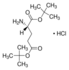 | 296 | -7.0 | -58.5 | -55.2 | -42.1 | -29.9 |
| 2 | F6 | 13033-84-6- | 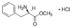 | 216 | -1.5 | -63.1 | -52.9 | -23.8 | -15.1 |
| 2 | F7 | H-p-iodo-DL-phe-OH | 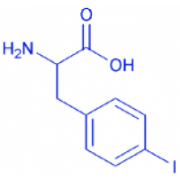 | 291 | -7.9 | -37.2 | -25.1 | -15.9 | -12.5 |
| 2 | F8 | 04-12-5074 | H-Glu(Obzl)-Obzl/p-Tosylate | 500 | -3.4 | -72.3 | -58.4 | -21.0 | -19.8 |
| 2 | F9 | H-Orn(Z)-Obzl | 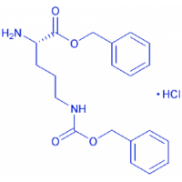 | 393 | -5.0 | -25.7 | -23.8 | -42.8 | -18.0 |
| 2 | F10 | H-Thr(tBu)otBuHCl | 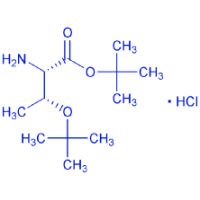 | 231 | -11.3 | -77.2 | -63.5 | -10.9 | -34.8 |
| 2 | G1 | 23239-35-2 | 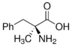 | 179 | -32.3 | -22.3 | -34.1 | -28.3 | -19.0 |
| 2 | G2 | 632-12-2 | 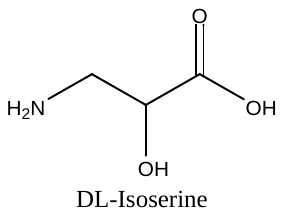 | 105 | -6.6 | -69.9 | -58.7 | -20.9 | -15.2 |
| 2 | G3 | 7389-87-9 | 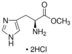 | 242 | 11.7 | -37.6 | -20.8 | -12.9 | -5.9 |
| 2 | G4 | 04-12-5004 | 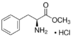 | 216 | -5.4 | -48.5 | -42.3 | -13.9 | -13.7 |
| 2 | G5 | H-D-Asp(Obzl)-OH | 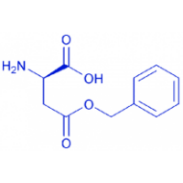 | 223 | -2.2 | -40.9 | -33.5 | -13.8 | -17.7 |
| 2 | G6 | 5856-62-2 | 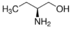 | 89 | -5.9 | -78.7 | -63.1 | -52.3 | -35.9 |
| 2 | G7 | 13472-00-9 | 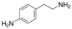 | 136 | -2.8 | -85.2 | -80.9 | -27.8 | -26.0 |
| 2 | G8 | 5856-63-3 | 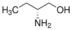 | 89 | 2.1 | -76.4 | -77.2 | -25.4 | -25.2 |
| 2 | G9 | 1492-24-6 | 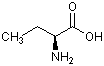 | 103 | 34.7 | -68.3 | -68.9 | -38.5 | -19.9 |
| 2 | G10 | L-alanylglycine | 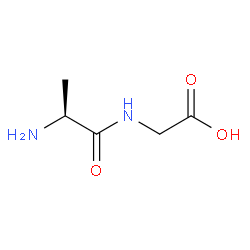 | 146 | -1.7 | -65.6 | -62.7 | -32.7 | -31.8 |
| 3 | A1 | 2432-74-8 | 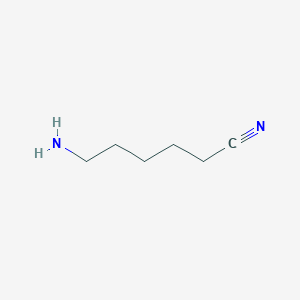 | 112 | 66.8 | -79.8 | -71.2 | -30.0 | -29.9 |
| 3 | A2 | 56.12.2 | 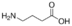 | 103 | 15.7 | -100.8 | -94.2 | -40.1 | -27.4 |
| 3 | A3 | 929-17-9 | 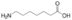 | 145 | -3.2 | -101.1 | -95.1 | -44.4 | -35.9 |
| 3 | A4 | 1187-42-4 | 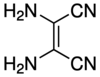 | 108 | 15.6 | -39.9 | -26.1 | -20.6 | -14.4 |
| 3 | A5 | 4-amino-1-butanol | 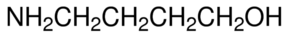 | 89 | 36.3 | -89.6 | -84.6 | -30.3 | -31.7 |
| 3 | A6 | 96-20-8 | 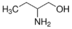 | 89 | -2.0 | -84.7 | -64.4 | -30.2 | -21.2 |
| 3 | A7 | 50910-54-8 | 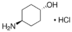 | 152 | -2.6 | -75.3 | -66.2 | -45.1 | -42.3 |
| 3 | A8 | L-cysteine | 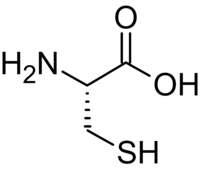 | 121 | -6.9 | -39.7 | -30.4 | -28.5 | -23.6 |
| 3 | A9 | [2-(aminomethyl)phenyl]acetic acid | 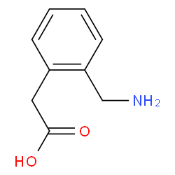 | 165 | -1.4 | -68.1 | -60.5 | -25.1 | -15.8 |
| 3 | A10 | 79286-79-6 | 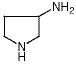 | 86 | -8.3 | -85.0 | -81.9 | -38.2 | -30.3 |
| 3 | B1 | see structure | 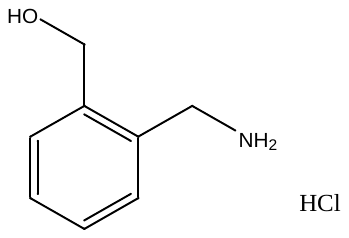 | 174 | 2.8 | -79.1 | -66.8 | -36.6 | -31.0 |
| 3 | B2 | 133437-08-8 | 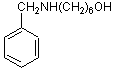 | 207 | -3.7 | -43.8 | -34.4 | -21.0 | -14.6 |
| 3 | B3 | 24123-14-6 | 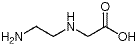 | 118 | -1.1 | -75.7 | -62.6 | -33.4 | -29.1 |
| 3 | B4 | 5098-14-6 | 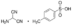 | 253 | -17.9 | -60.0 | -38.1 | -23.3 | -17.9 |
| 3 | B5 | 2079-89-2 | 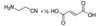 | 128 | -6.4 | -73.7 | -66.5 | -33.8 | -33.9 |
| 3 | B6 | 1002-57-9 | 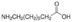 | 159 | -2.0 | -96.3 | -85.8 | -41.6 | -33.8 |
| 3 | B7 | 1-benzyl-3-(methylamino)pyrrolidine | 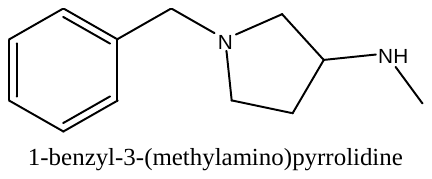 | 190 | -12.2 | -50.3 | -42.7 | -21.5 | -16.9 |
| 3 | B8 | 693-57-2 | 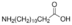 | 215 | -5.8 | -36.6 | -32.0 | -9.1 | -14.1 |
| 3 | B9 | 17702-88-4 | 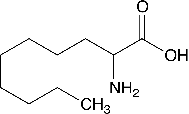 | 187 | -21.8 | -51.1 | -42.9 | -28.9 | -20.8 |
| 3 | B10 | 23159-07-1 | 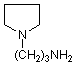 | 128 | -3.0 | -91.3 | -89.0 | -25.6 | -16.2 |
| 3 | C1 | 1197-18-8 | 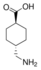 | 157 | 7.8 | -111.2 | -103.3 | -33.5 | -35.7 |
| 3 | C2 | 163061-73-2 | 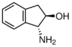 | 149 | -1.7 | -57.0 | -44.1 | -26.2 | -23.5 |
| 3 | C3 | 107-95-9 | 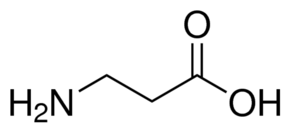 | 89 | 2.1 | -87.0 | -75.3 | -48.0 | -35.6 |
| 3 | C4 | 5036-48-6 |  | 125 | 46.5 | -86.7 | -71.3 | -29.7 | -23.1 |
| 3 | C5 | 108-91-8 |  | 99 | 1.1 | -76.4 | -63.1 | -29.9 | -23.3 |
| 3 | C6 | 768-94-5 |  | 151 | 3.3 | -44.4 | -30.7 | -20.9 | -21.7 |

|  |  |  |  | Inhibition %  Amine only | Meta Inhibition %  2µM final rep1 | Meta Inhibition %  2µM final rep2 | Para Inhibition %  2µM final rep1 | Para Inhibition %  2µM final rep2 |
| --- | --- | --- | --- | --- | --- | --- | --- | --- |
| A1 | 5382-16-1 |  | 101 | -4.7 | -61.6 | -61.8 | -16.0 | -11.2 |
| A2 | 16652-71-4 |  | 241.7 | 14.3 | -61.6 | -46.9 | -20.2 | -17.0 |
| A3 | 36520-39-5 |  | 93.56 | -23.7 | -74.7 | -58.7 | -30.7 | -27.2 |
| A4 | 6921-28-4 |  | 93.13 | 6.7 | -46.0 | -68.5 | -15.5 | -10.7 |
| A5 | 111-95-5 |  | 133 | 68.6 | -43.2 | -37.7 | -16.7 | -13.2 |
| A6 | (1R,2S,5R,6S)-9-azabicyclo[4.2.1]nonane-2,5-diol |  | 157 | 46.4 | -65.1 | -59.2 | -27.5 | -21.3 |
| A7 | 626-56-2 |  | 99.18 | 47.5 | -46.3 | -45.1 | -21.5 | -19.4 |
| A8 | 109-01-3 |  | 100.16 | 5.5 | -57.7 | -51.5 | -27.3 | -16.9 |
| A9 | 18621-18-6 |  | 109.56 | -6.3 | -55.9 | -45.2 | -19.4 | -11.8 |
| A10 | 16369-21-4 |  | 103.17 | 3.3 | -55.7 | -49.1 | -21.6 | -20.9 |
| B1 | 101-83-7 |  | 181 | -4.7 | -54.0 | -43.5 | -26.3 | -27.7 |
| B2 | 60399-02-2 |  | 173.64 | 4.9 | -42.5 | -31.5 | -8.8 | -13.3 |
| B3 | 7755-92-2 |  | 114.15 | -1.4 | -48.2 | -42.7 | -17.4 | -9.1 |
| B4 | 109-01-3 |  | 100 | -7.6 | -59.1 | -43.6 | -24.2 | -21.5 |
| B5 | 111-42-2 |  | 105 | 6.0 | -48.8 | -39.1 | -22.5 | -15.9 |
| B6 | 172603-05-3 |  | 200 | 3.5 | -48.5 | -34.3 | -10.3 | -7.1 |
| B7 | 6511-88-2 |  | 181 | -6.0 | -58.8 | -45.8 | -23.6 | -21.4 |
| B8 | 99724-19-3 |  | 186 | 56.5 | -65.7 | -58.3 | -26.0 | -16.8 |
| B9 | 745048-12-8 |  | 155 | -11.3 | -69.0 | -65.0 | -27.0 | -27.5 |
| B10 | 4,4-diethoxypiperidine hydrochloride |  | 173 | 5.4 | -67.5 | -64.2 | -22.6 | -20.0 |
| C1 | (S)-2-(azidomethyl)-1,4-dioxa-8-azaspiro[4.5]decane |  | 198.11 | 17.4 | -49.3 | -39.0 | 6.6 | 11.0 |
| C2 | 35161-71-8 |  | 69.11 | 6.1 | -49.9 | -40.2 | -17.1 | -8.4 |
| C3 |  |  |  | 8.9 | -46.6 | -33.2 | -9.6 | -7.9 |
| C4 |  |  |  | 0.6 | -40.8 | -21.5 | 23.8 | -13.2 |
| C5 |  |  |  | 14.2 | -56.1 | -45.0 | -80.1 | -24.4 |
| C6 | morpholine |  | 87 | 16.6 | -37.9 | -7.9 | -2.9 | 0.7 |
| C7 | 4-Methylpiperidine |  | 99 | -1.5 | -52.5 | -41.7 | -12.7 | -6.5 |
| C8 | dimethyl amine |  | 45 | -0.1 | -44.4 | -28.6 | -8.8 | 9.2 |
| C9 | diethylamine |  | 73 | -4.5 | -48.2 | -36.4 | -16.7 | -5.3 |
| C10 | pyrrolidine |  | 71 | 7.6 | -60.9 | -60.6 | -18.9 | -19.9 |
| D1 | 51-35-4 |  | 131.13 | 2.1 | -77.6 | -66.9 | -16.5 | -12.6 |
| D2 | 609-36-9 |  | 115.13 | 10.2 | -64.9 | -54.9 | -17.7 | -10.6 |
| D3 | 344-25-2 |  | 115 | 1.2 | -78.1 | -71.0 | -14.6 | -12.1 |
| D4 | 110-85-0 |  | 86 | 5.1 | -54.4 | -34.9 | 0.4 | -14.0 |
| D5 | 768-66-1 |  | 141.25 | -0.5 | -50.5 | -42.6 | -19.8 | -19.2 |
| D6 | 2812-46-6 |  | 171.24 | -5.6 | -46.7 | -40.6 | -19.9 | -16.3 |
| D7 | 51207-66-0 |  | 154.25 | -5.6 | -71.6 | -42.9 | -20.4 | -14.4 |
| D8 | 1484-84-0 |  | 129.2 | -7.6 | -46.0 | -37.8 | -21.4 | -17.4 |
| D9 | 177-11-7 |  | 143.18 | 2.8 | -61.5 | -50.3 | -13.6 | -7.8 |
| D10 | 169447-86-3 |  | 276.38 | -5.2 | -45.5 | -38.3 | -25.8 | -20.5 |
| E1 | 2328.12.3 |  | 229.7 | 49.1 | -40.5 | -28.0 | -2.3 | 1.0 |
| E2 | 622-26-4 |  | 129.2 | 21.0 | -36.4 | -52.9 | 17.7 | 19.3 |
| E3 | 1683-49-4 |  | 282 | 2.6 | -41.9 | -31.7 | 2.4 | 7.0 |
| E4 | 31252-42-3 |  | 175 | -1.3 | -37.3 | -36.5 | -6.1 | -5.5 |
| E5 | 39546-32-2 |  | 128 | -0.7 | -70.5 | -58.7 | -19.9 | -14.7 |
| E6 | 57988-58-6 |  | 256 | 12.1 | -13.6 | 2.2 | 3.3 | 11.8 |
| E7 | 196204-01-0 |  | 205 | 42.6 | -82.6 | -66.2 | -24.5 | -20.5 |
| E8 | 1153950-54-9 |  | 132.59 | 62.2 | -70.6 | -58.8 | -16.4 | -15.5 |
| E9 | 1153950-49-2 |  | 132.59 | 0.9 | -64.2 | -55.3 | -14.0 | -7.8 |
| E10 | 136725-53-6 |  | 126 | 1.7 | -44.0 | -65.1 | -19.1 | -14.3 |
| F1 | 6000-50-6 |  | 193 | -0.8 | -52.5 | -53.1 | -20.0 | -10.7 |
| F2 | 132958-72-6 |  | 114 | -3.7 | -75.8 | -67.8 | -32.7 | -16.9 |
| F3 | 132883-44-4 |  | 114 | -9.9 | -74.6 | -63.6 | -24.8 | -9.3 |
| F4 | 63468-63-3 |  | 105.57 | 16.0 | -70.0 | -61.7 | -24.0 | -17.4 |
| F5 | 147740-02-1 |  | 193.1 | 6.6 | -39.0 | -33.8 | -21.5 | -11.2 |
| F6 | (1S,2R)-1,2-diphenyl-2-(piperazin-1-yl)ethan-1-ol |  | 282 | 4.4 | -34.3 | -25.5 | -10.3 | -0.9 |
| F7 | ethyl 3-phenylaziridine-2-carboxylate |  | 191 | 34.2 | -49.9 | -38.9 | -19.4 | -14.9 |
| F8 | 4-nitro-N-(1-phenoxy-3-(piperazin-1-yl)propan-2-yl)benzenesulfonamide |  | 420 | 57.1 | -27.6 | -17.2 | -11.6 | -46.1 |
| F9 | 1-((1R,2S)-1,2-diphenyl-2-(piperazin-1-yl)ethyl)-3-propylurea |  | 367 | 51.3 | -12.2 | -18.3 | -0.1 | 1.6 |
| F10 |  |  |  | 2.0 | -47.8 | -44.6 | -27.1 | -25.9 |
| G1 | 1121-92-2 |  | 113.2 | -8.5 | -61.0 | -45.7 | -27.8 | -22.9 |
| G2 | 11-49-9 |  | 99.17 | -5.8 | -63.4 | -51.9 | -23.9 | -3.4 |
| G3 | 505-19-1 |  | 86.14 | 6.4 | -40.4 | -32.9 | -24.6 | -19.6 |
| G4 | 1126-09-6 |  | 157 | 6.0 | -59.9 | -48.1 | -23.5 | -24.5 |
| G5 | 1135-40-6 |  | 221 | 8.5 | -48.5 | -27.9 | -17.3 | -14.3 |
| G6 | 29915-38-6 |  | 243 | 200.9 | -50.2 | -41.3 | -26.1 | -23.5 |
| G7 | 7365-44-8 |  | 229 | 0.1 | -44.4 | -37.2 | -24.6 | -15.8 |
| G8 | 68399-81-5 |  | 259.28 | -9.2 | -50.0 | -41.0 | -24.8 | -25.1 |
| G9 | 7365-82-4 |  | 182 | -3.4 | -47.7 | -53.0 | -23.0 | -13.2 |
| G10 | 165528-81-4 |  | 228 | 0.5 | -43.9 | -34.9 | -8.9 | -6.6 |
