## Supplemental Table 3 for "Chemical inhibition of ENL/AF9 YEATS domains in acute leukemia"

| plate# | well | CAS | Structure | MW | IC50, M |
| --- | --- | --- | --- | --- | --- |
| 1 | A2 | 929-06-6 | 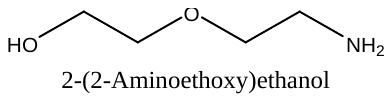 | 105.14 | 5.49E-07 |
| 1 | A4 | 13325-10-5 | 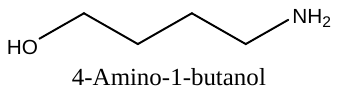 | 89 | 4.72E-07 |
| 1 | A8 | 156-87-6 | 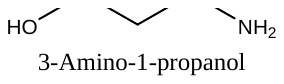 | 75 | 5.63E-07 |
| 1 | B9 | 123-00-2 | 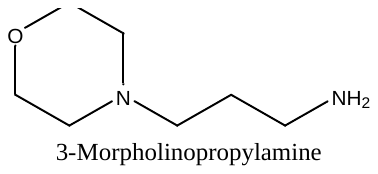 | 144 | 5.52E-07 |
| 1 | C4 | 56-92-8 | 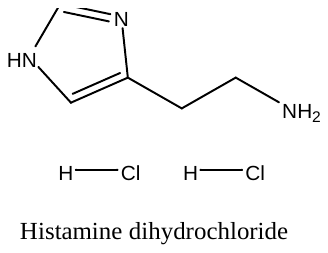 | 184 | 1.22E-07 |
| 1 | C5 | 696-60-6 | 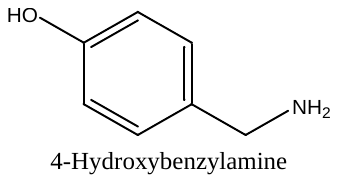 | 123 | 3.71E-07 |
| 1 | C10 | 35303-76-5 | 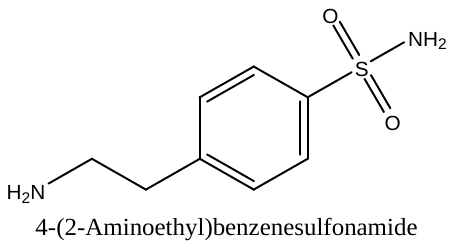 | 200 | 5.15E-07 |
| 1 | F7 | 30433-91-1 | 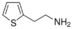 | 127 | 1.36E-06 |
| 1 | G1 | 100-82-3 | 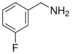 | 125 | 5.19E-07 |
| 1 | G5 | 140-75-0 | 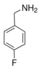 | 125 | 1.02E-06 |
| 1 | G7 | 593-51-1 | 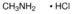 | 68 | 3.88E-07 |
| 2 | A9 | 7663-77-6 | 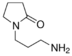 | 142 | 8.76E-07 |
| 2 | D8 | 14907-27-8 | 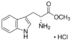 | 255 | 8.73E-07 |
| 2 | D9 | 04-13-5043 | 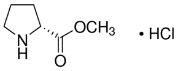 | 166 | 9.50E-07 |
| 2 | E3 | 04-12-5044 | 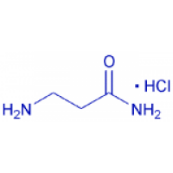 | 125 | 5.18E-07 |
| 2 | E8 | 141-43-5 | 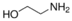 | 61 | 4.26E-07 |
| 2 | G7 | 13472-00-9 | 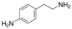 | 136 | 3.17E-07 |
| 3 | A1 | 2432-74-8 | 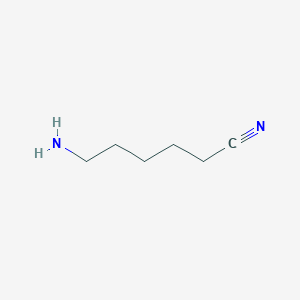 | 112 | 5.54E-07 |
| 3 | A2 | 56.12.2 | 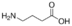 | 103 | 6.93E-07 |
| 3 | A3 | 929-17-9 | 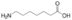 | 145 | 3.61E-07 |
| 3 | A5 | 4-amino-1-butanol | 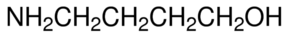 | 89 | 2.98E-07 |
| 3 | A10 | 79286-79-6 | 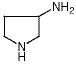 | 86 | 2.30E-07 |
| 3 | B6 | 1002-57-9 | 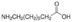 | 159 | 4.55E-07 |
| 3 | B10 | 23159-07-1 | 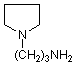 | 128 | 3.36E-07 |
| 3 | C1 | 1197-18-8 | 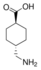 | 157 | 3.24E-07 |
| 3 | C3 | 107-95-9 | 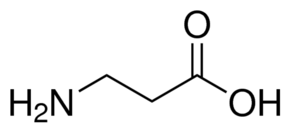 | 89 | 6.27E-07 |
| 3 | C4 | 5036-48-6 | 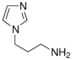 | 125 | 3.31E-07 |
|  | D3 | 344-25-2 | 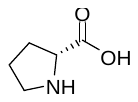 | 115 | 6.11E-07 |
