## Supplemental Table 4 for "Chemical inhibition of ENL/AF9 YEATS domains in acute leukemia"

| # | well | CAS | Structure | MW | Control rep1 | Rep2 | Rep3 | Rep4 | SuFEx rep1 | Rep2 | Rep3 | Rep4 |
| --- | --- | --- | --- | --- | --- | --- | --- | --- | --- | --- | --- | --- |
| 1 | A1 | 153-98-0 | 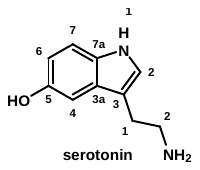 | 176 | 102 | 70 | 83 | 73 | 133 | 120 | 139 | 123 |
| 2 | A2 | 61-54-1 | 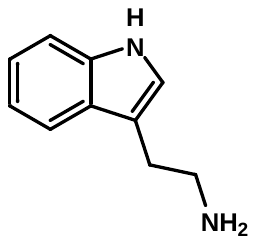 | 160 | 104 | 81 | 92 | 84 | 142 | 124 | 138 | 131 |
| 3 | A3 | 617-89-0 | 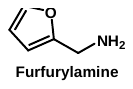 | 97 | 144 | 100 | 115 | 106 | 171 | 166 | 185 | 170 |
| 4 | A4 | 5036-48-6 | 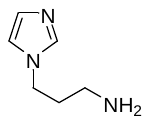 | 125 | 156 | 125 | 130 | 116 | 142 | 139 | 162 | 151 |
| 5 | A5 | 51387-90-7 | 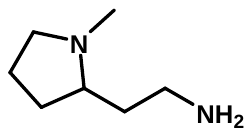 | 128 | 155 | 115 | 118 | 108 | 176 | 152 | 194 | 85 |
| 6 | A6 | 7154-73-6 | 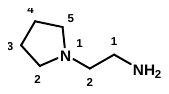 | 114 | 120 | 92 | 110 | 107 | 122 | 115 | 138 | 114 |
| 7 | A7 | 696-60-6 | 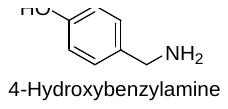 | 123 | 120 | 99 | 121 | 103 | 140 | 116 | 151 | 143 |
| 8 | A8 | 3529-08-6 | 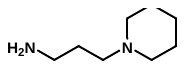 | 142 | 124 | 82 | 115 | 81 | 173 | 162 | 225 | 183 |
| 9 | A9 | 140-31-8 | 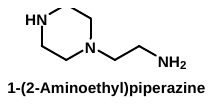 | 129 | 108 | 89 | 117 | 105 | 127 | 119 | 138 | 133 |
| 10 | A10 | 3399-67-5 | 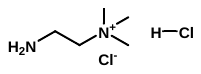 | 175.1 | 115 | 100 | 112 | 101 | 144 | 132 | 89 | 77 |
| 11 | B1 | 100-82-3 | 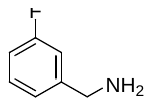 | 125 | 108 | 100 | 116 | 100 | 135 | 123 | 138 | 129 |
| 12 | B2 | 123-00-2 | 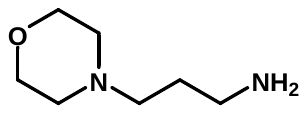 | 144 | 90 | 98 | 108 | 98 | 147 | 137 | 146 | 149 |
| 13 | B3 | 81028-90-2 | 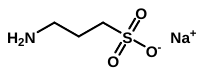 | 161 | 102 | 76 | 89 | 80 | 105 | 72 | 86 | 87 |
| 14 | B4 | 4795-29-3 | 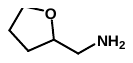 | 101 | 114 | 95 | 102 | 112 | 169 | 113 | 132 | 140 |
| 15 | B5 | 13258-63-4 | 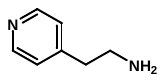 | 122 | 96 | 71 | 95 | 80 | 149 | 76 | 99 | 93 |
| 16 | B6 | 19968-85-5 | 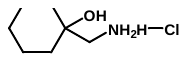 | 166 | 128 | 91 | 117 | 105 | 94 | 107 | 99 | 80 |
| 17 | B7 | 156-87-6 | 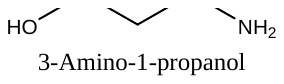 | 75 | 113 | 92 | 117 | 98 | 97 | 76 | 142 | 80 |
| 18 | B8 | 534-03-2 | 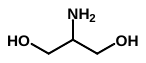 | 91 | 108 | 94 | 130 | 97 | 97 | 74 | 90 | 73 |
| 19 | B9 | 1115-90-8 | 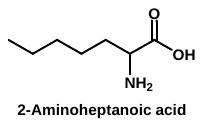 | 145 | 114 | 94 | 124 | 99 | 97 | 113 | 148 | 117 |
| 20 | B10 | 138060-07-8 | 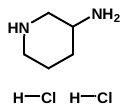 | 173 | 109 | 96 | 126 | 100 | 153 | 138 | 148 | 127 |
| 21 | C1 | 79467-22-4 | 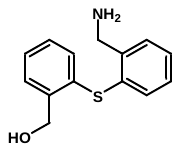 | 245 | 121 | 108 | 128 | 110 | 99 | 82 | 89 | 81 |
| 22 | C2 | 15996-76-6 | 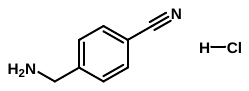 | 168.6 | 97 | 80 | 109 | 74 | 102 | 107 | 90 | 78 |
| 23 | C3 | 22287-35-0 | 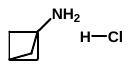 | 120 | 137 | 76 | 111 | 100 | 153 | 124 | 150 | 123 |
| 24 | C4 | 6530/09/2 | 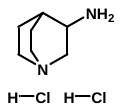 | 199 | 137 | 104 | 109 | 106 | 165 | 153 | 168 | 164 |
| 25 | C5 | 6246-48-6 | 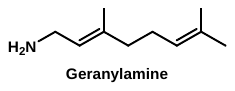 | 153.27 | 107 | 105 | 118 | 97 | 89 | 85 | 94 | 91 |
| 26 | C6 | 57260-73-8 | 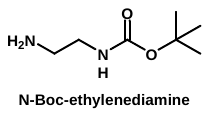 | 160 | 110 | 99 | 111 | 113 | 149 | 133 | 161 | 143 |
| 27 | C7 |  | 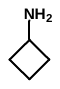 | 71 | 124 | 97 | 109 | 107 | 149 | 153 | 161 | 87 |
| 28 | C8 | cds000177-250 | 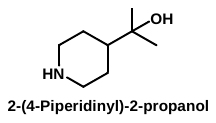 | 143 | 105 | 94 | 119 | 104 | 108 | 83 | 95 | 75 |
| 29 | C9 | 334618-07-4 | 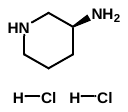 | 173 | 93 | 83 | 106 | 90 | 84 | 145 | 169 | 89 |
| 30 | C10 | 334618-23-4 | 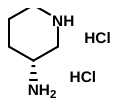 | 173 | 111 | 97 | 91 | 100 | 139 | 145 | 182 | 165 |
| 31 | D1 | 35386-24-4 | 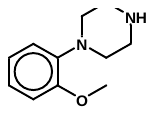 | 192 | 97 | 98 | 92 | 82 | 104 | 114 | 124 | 113 |
| 32 | D2 | 4403-69-4 | 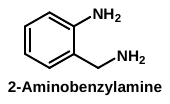 | 122.17 | 111 | 99 | 104 | 93 | 124 | 138 | 154 | 148 |
| 33 | D3 | 4403-70-7 | 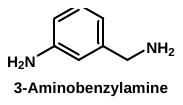 | 122 | 93 | 84 | 98 | 77 | 81 | 91 | 147 | 149 |
| 34 | D4 | 307496-23-7 | 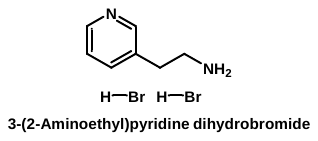 | 283.99 | 112 | 96 | 119 | 102 | 138 | 132 | 149 | 142 |
| 35 | D5 | 13325-10-5 | 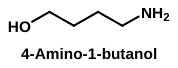 | 89 | 119 | 84 | 105 | 100 | 91 | 71 | 79 | 90 |
| 36 | D6 | 56-92-8 | 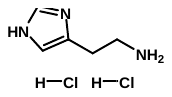 | 184 | 133 | 76 | 112 | 115 | 157 | 69 | 86 | 92 |
| 37 | D7 | 123-00-2 | 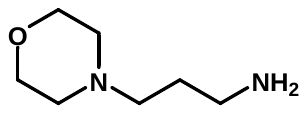 | 144 | 95 | 76 | 95 | 95 | 158 | 70 | 136 | 83 |
| 38 | D8 | 27578-60-5 | 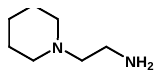 | 128 | 111 | 87 | 121 | 113 | 176 | 158 | 197 | 207 |
| 39 | D9 | 1197-18-8 | 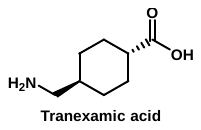 | 157 | 114 | 90 | 108 | 100 | 94 | 74 | 100 | 88 |
| 40 | D10 | 2432-74-8 | 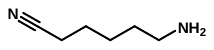 | 112 | 110 | 97 | 117 | 84 | 142 | 125 | 155 | 137 |
| 41 | E1 | 13472-00-9 | 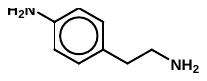 | 136 | 125 | 83 | 119 | 101 | 162 | 153 | 188 | 167 |
| 42 | E2 | 1002-57-9 | 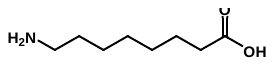 | 159 | 97 | 79 | 102 | 85 | 173 | 159 | 200 | 175 |
| 43 | E3 | 929-17-9 | 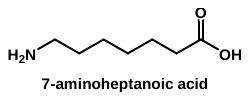 | 145 | 92 | 79 | 94 | 80 | 160 | 142 | 190 | 184 |
